# Dosage compensation and meiotic sex chromosome inactivation are maintained in the absence of selection

**DOI:** 10.1101/2025.07.27.667025

**Authors:** Darren J. Parker, Zoé Dumas, Rocío Gómez Lencero, Jean-Marc Aury, Marjorie Labedan, Patrick Tran Van, Benjamin Istace, Corinne Cruaud, Karine Labadie, Benjamin Noel, Susana Freitas, Jelisaveta Djordjevic, Tanja Schwander

## Abstract

Dosage compensation and meiotic sex chromosome inactivation (MSCI) are key mechanisms regulating gene expression from the X chromosome in male-heterogametic species. While the convergent evolution of these mechanisms is well-documented, their evolutionary fate under relaxed selection remains poorly understood. Here, we test whether dosage compensation and MSCI persist following three independent transitions to parthenogenesis in stick insects, where selection on male phenotypes is relaxed. Using rare males occasionally produced by parthenogenetic females, chromosome-level genome assemblies, RNA-seq from multiple tissues, and immunocytochemistry, we find that dosage compensation is fully conserved across all nine studied somatic tissues. This is even the case in the oldest, approximately 1.5 million years old all-female lineage, and for tissue-specific genes for which dosage variation is not expected to be very deleterious. Meiotic X inactivation in the germline is also conserved. Surprisingly however, expression data and cytological markers indicate that MSCI signatures are even stronger in parthenogenetic males, a pattern likely driven by prolonged autosomal transcription during meiosis. These results indicate that X-targeting dosage compensation and MSCI are highly stable over evolutionary time and may be maintained in all-female lineages by a combination of evolutionary constraint, pleiotropy, or very weak selection, whereas autosomal expression during meiosis shifts rapidly under relaxed selection.

**Significance Statement:** Sex chromosomes are regulated by specialized mechanisms that balance gene expression between males and females in somatic tissues (dosage compensation) and silence the X chromosome during male meiosis (meiotic sex chromosome inactivation, MSCI). These processes are thought to decay when they become redundant, yet suitable systems to test this prediction are scarce. We studied rare males from parthenogenetic stick insect lineages that have been evolving without sex for up to 1.5 million years. Surprisingly, both dosage compensation and MSCI remain fully functional, despite relaxed selection. Instead, we find altered autosomal expression during meiosis. Our results reveal that sex chromosome regulation is evolutionarily stable and constrained, persisting even after the loss of its purpose.

## Introduction

Sexual reproduction is key to the diversity of life and the main driver of species’ lifecycles. In eukaryotic species with distinct male and female sexes, sex determination is frequently governed by sex chromosomes such as X and Y chromosomes in male heterogametic species (1). Because males carry only a single X chromosome while females carry two, X chromosome expression is regulated by highly specialized mechanisms: dosage compensation (in somatic cells) and meiotic sex chromosome inactivation (MSCI, in germ cells). Dosage compensation is a mechanism that balances the expression of X-linked and autosomal genes independently of the number of X chromosome copies. It has evolved independently across a wide range of taxa (2–4) including mammals (5), reptiles (6), fish (7, 8), nematodes (9), and insects (10–16). However, the X is typically not dosage compensated in gonads, where expression from the sex chromosome(s) is further often suppressed via MSCI (2, 12, 13, 17, 18). Specifically, MSCI involves the inactivation of heteromorphic sex chromosomes during meiosis through the addition of repressive chromatin marks, likely due to the inability of the X and the degraded (or completely lacking) Y to synapse at the onset of meiosis (19).

While far from ubiquitous (4, 20), dosage compensation and MSCI are typically maintained across clades for millions of years. Furthermore, a breadth of studies on both mechanisms indicate that they can evolve convergently as new sex chromosomes arise, involving similar or different mechanisms. This is revealed for example by research in dipterans, where X chromosomes have evolved independently from ancestral autosomes, with different dosage compensation mechanisms emerging in parallel (21–28). Similarly, the independent evolution of MSCI mechanisms in divergent taxa involves both shared and lineage-specific components (e.g. (29–31)).

In contrast to the evidence for the convergent *de novo* evolution of dosage compensation and MSCI, it remains unknown how ancestral dosage compensation and MSCI systems evolve once they become redundant, which would be the case upon the evolution of a novel sex chromosome system. This is in part because suitable model systems with sex chromosome turnovers and accompanying evolution of X-targeting regulatory mechanisms remain understudied. Furthermore, sex chromosome turnover events in species with established dosage compensation and MSCI systems are expected to be rare because the maintenance of active systems could cause extensive misregulation of gene expression on the newly-formed sex chromosomes (32–34). A unique opportunity to overcome such constraints is to study the fate and potential decay of dosage compensation and MSCI in all-female parthenogenetic species. Following a transition to parthenogenesis, the X chromosome effectively adopts an autosomal inheritance pattern as it no longer varies in copy numbers since all individuals are female and carry two X copies. As a result, if dosage compensation or MSCI is costly (because of the cost of the machinery or due to maladaptive expression in females) we would expect selection to reduce a species’ ability to perform these mechanisms following a transition to parthenogenesis. In the absence of such costs in females, dosage compensation and MSCI would be expected to decay and vestigialize as a consequence of drift. A reduction of the two X regulatory mechanisms is difficult to observe in parthenogenetic species as we need to examine expression in both sexes. This barrier can be overcome in some systems if rare males are produced, a phenomenon frequently observed in parthenogenetic arthropods (35). The production of rare males would be impossible for species where the perturbation of dosage compensation is lethal in males, as is the case for certain mutations in the *Drosophila* fly dosage compensation machinery (e.g., (36)). The suppression of dosage compensation does however not affect male viability in *Anopheles* mosquitoes, and instead causes a developmental delay (25). This variation between taxa illustrates that inference regarding the functioning and maintenance of dosage compensation cannot be based on the observation of male phenotypes in parthenogenetic species. With respect to X regulation in the germline, X-autosome translocations negatively affect fertility in *Drosophila* flies (37, 38), where X regulation happens via a mechanism that differs from classical MSCI as described in mammals and other insect taxa (17, 39). For other insects, fertility consequences of MSCI perturbation are unknown. As a consequence, fertility or sperm viability phenotypes in rare males do not provide information about the functioning of MSCI.

Here we take advantage of such rare males to examine dosage compensation and MSCI in three species of parthenogenetic *Timema* stick insects, using a combination of new chromosome-level genome assemblies, RNA sequencing of eight different tissues, and cell biology. *Timema* have had multiple independent transitions to parthenogenetic reproduction over the last 0.5-1.5 million years (40–43). Previously we have found that sexual *Timema* species show complete dosage compensation in somatic tissues throughout development, and MSCI in the male germline (11, 17, 44). This targeted regulation of the X chromosome likely involves general molecular mechanisms shared across insects (17) given that the stick insect X chromosome likely evolved prior to the diversification of insects over 450 mya (33). We can examine changes in dosage compensation and MSCI in parthenogenetic *Timema* as they occasionally produce rare males via two different processes. Some males arise from aneuploidy of the X chromosome (45), which since *Timema* have an XX/X0 sex-determination system, results in the production of a male. Others develop as a consequence of very rare instances of sex within parthenogenetic populations (46). Field and experimental evidence indicates that males produced by parthenogenetic *Timema* are exceedingly rare and exhibit reduced reproductive performance (45). During extensive field collections conducted over multiple years (2007–2012), only 11 males were recovered across several independently derived asexual lineages, despite the simultaneous collection of very large numbers of females. Functional assays of sperm performance further demonstrate that while rare males are capable of fertilization when mated as the sole partner of virgin females from closely related sexual species, reproductive success is low and they fail entirely under sperm competition (45). Thus, male production in parthenogenetic *Timema* is both extremely infrequent and associated with reduced reproductive performance. These features indicate relaxed selection on male-specific traits, while still allowing the occasional availability of males for mechanistic analyses of dosage compensation and meiotic sex chromosome inactivation. Contrary to our expectations we find that the targeted regulation of the X chromosome underlying dosage compensation and MSCI has been fully conserved in each of the three parthenogenetic lineages, with relaxed selection affecting autosomal rather than X-linked expression in male meiotic cells. These results indicate that selection against X-regulatory mechanisms is weak or that changes to dosage compensation and MSCI mechanisms are highly constrained.

## Results and Discussion

### Genome assemblies reveal a highly conserved X chromosome contrasted by autosomal fissions and fusions

We generated chromosome-level genome assemblies for two of the three sexual species of *Timema* used in this study, *T. cristinae* and *T. podura*, to complement the available assembly of *T. poppense* (Table S1). In both new assemblies, the number of large scaffolds assembled (hereafter referred to as chromosomes) matched previous karyotype information (40) and coverage of male-vs female-derived short reads were used to identify the X chromosome (Supplemental Info). Annotation using a combination of gene predictions, protein, and RNAseq evidence from each species and its parthenogenetic sister species and inference of 8118 one-to-one orthologs (see Methods) revealed that the X chromosome content has not changed in any of the sexual or asexual species, as expected given the long-term conservation of X chromosome content in stick insects (11, 47). Using comparisons with a stick insect (*Bacillus rossius*) from the Euphasmatodea clade (which split over 120 million years ago from *Timema* (48), we further find that even X chromosome synteny is highly conserved, while many rearrangements are apparent on the autosomes (Fig. 1). Specifically, the change in karyotype from 14 to 12 chromosomes during *Timema* evolution (40) involved at least two fissions and three fusions involving *T. podura* chromosomes 2, 3, 5 and 10 (Fig. 1, S1).

**Fig. 1:**
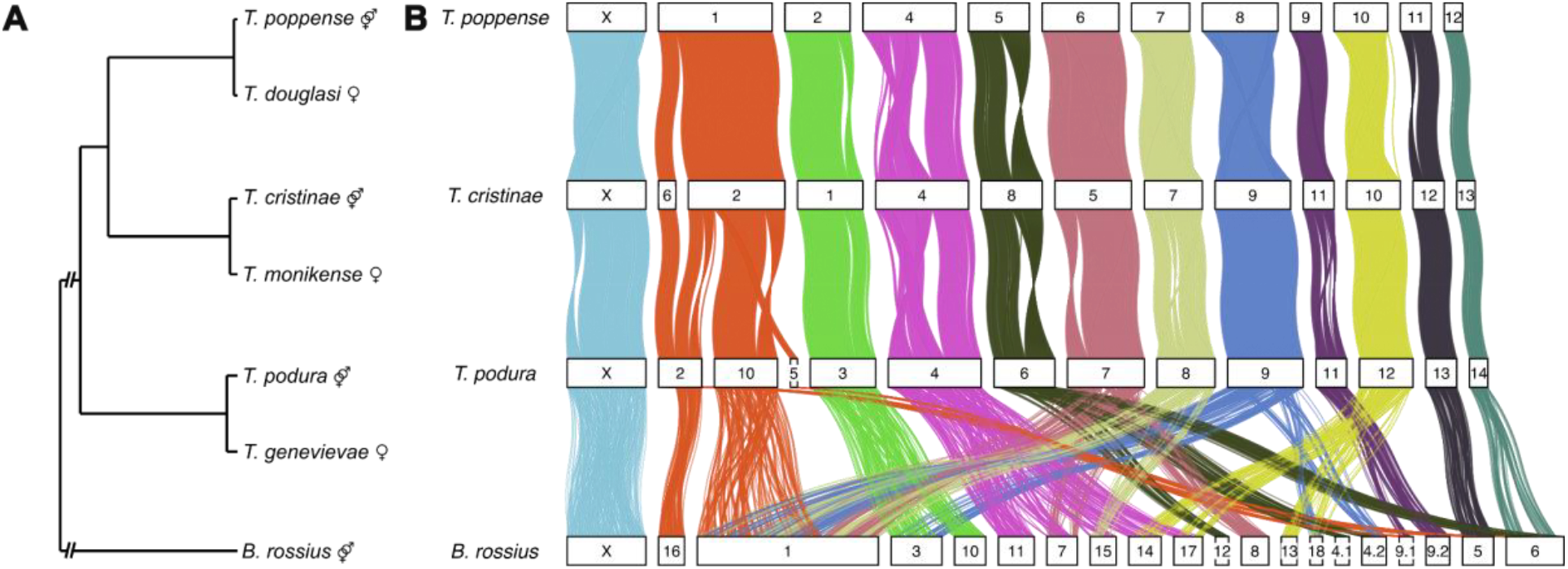
Autosomal genomic instability contrasts with X chromosome conservation. **A**. Simplified phylogeny of the used species, highlighting the independently evolved parthenogenetic *Timema* species and their closest sexual relatives. *Timema* are the sister clade to all other stick insects (including B. *rossius*), and branches reflecting this split have been shortened for display. B. Chord diagram showing synteny between the assemblies of sexual species, based on 4241 one-to-one orthologs across the focal *Timema* genomes and a distantly related stick insect (*Bacillus rossius*). See Fig. S1 for synteny based on 8118 one-to-one orthologs across *Timema* only.

### Dosage compensation in somatic tissues is not reduced in parthenogenetic species

Dosage compensation between the sexes in parthenogenetic species was very similar to that observed in sexual species. In all seven non-reproductive tissues (antennae, brain, defence glands, gut, fat body, femur, and tarsi) we observe almost complete dosage compensation in all species (Fig. 2). As expected, we observe some variation in the expression of X-linked genes between the sexes for specific tissues and species (resulting in small but significant differences in expression of autosomal versus X-linked genes). Such variation is typical in studies of dosage compensation (e.g., (13, 49, 50) and can be interpreted as sex-specific regulation of individual genes rather than a consequence of variable or incomplete dosage compensation. If dosage compensation was reduced in parthenogenetic species, we would expect to see a larger deviation from complete dosage compensation in parthenogenetic species and an accompanying lower male-to-female expression ratio for the X in parthenogenetic species when compared to the sexual sister species for tissues with almost complete dosage compensation. We do not observe this, instead finding only small variations in X-linked expression with no consistent pattern (Fig. 2, S2-S11, Table S2), showing that dosage compensation is maintained across tissues even under relaxed selection for up to 1.5 million years. This conclusion extends to more in depth analyses: the amount of dosage compensation is consistent between the sexual and parthenogenetic species on a gene-by-gene basis (Fig. S2-S4), and we also find no evidence for decreased efficiency of dosage compensation for genes with tissue-specific expression (Fig. S5-S8), which are expected to be relatively less constrained than housekeeping genes (51).

**Fig. 2:**
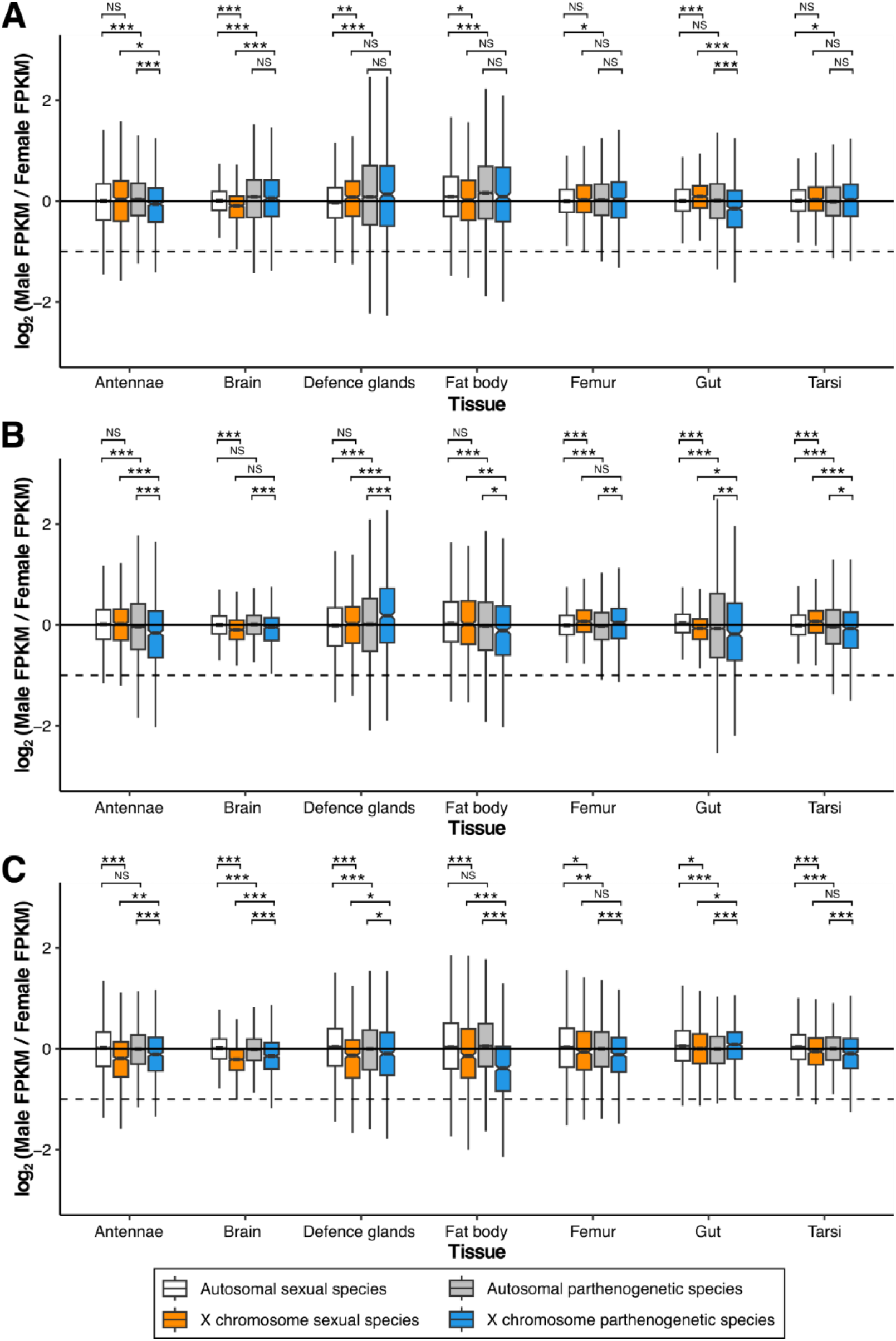
Male to female expression ratios for the X and autosomes in non-reproductive tissues reveal the maintenance of dosage compensation under relaxed selection. Dashed lines represent a two-fold reduction in expression in males (as expected if there was no dosage compensation). A. Sexual species = *T. cristinae*, parthenogenetic species = *T. monikense*. B. Sexual species = *T. podura*, parthenogenetic species = *T. genevievae*. C. Sexual species = *T. poppense*, parthenogenetic species = *T. douglasi*. Asterisks indicate the significance (FDR) of Wilcoxon tests (***<0.001, **<0.01, *<0.05, NS > 0.05).

Parthenogenetic species no longer require sex-specific regulation of X chromosome copy number across individuals. Under such conditions, the strength of selection acting on dosage compensation mechanisms is expected to be substantially reduced. Whether this relaxation should lead to detectable evolutionary decay depends on several unknown parameters, including the number of loci involved, mutational target size, and the degree of pleiotropy. If the machinery is costly or maladaptive in females, selection could favor its reduction. Alternatively, if costs are negligible, its evolution would be governed largely by drift. The absence of detectable change over up to 1.5 million years therefore indicates either unusually slow neutral decay, functional constraint, or the action of weak purifying selection, but does not allow us to distinguish conclusively among these alternatives. Support for functional constraints because of pleiotropic effects in females stems from evidence for the maintenance of dosage compensation-like mechanisms targeting the ancestral X in *Drosophila melanogaster* fruit flies (52) even tens of myr after a sex chromosome turnover, which reverted the ancestral X (now referred to as the dot) back to an autosome (15). Similar to our present study in parthenogenetic species, such reversals should result in relaxed selection for dosage compensation on the ancestral X (dot) chromosome. Yet, the dot is still regulated in a chromosome-specific manner, distinct from the other autosomes and the novel X chromosome. Thus, the dot is regulated via the protein POF (52), which might reflect vestiges of the former dosage compensation machinery when the dot was still the X chromosome.

A second reason to expect changes in dosage compensation could be if dosage compensation mechanisms result in maladaptive gene expression in females, i.e., partially unresolved sexual antagonism (53). This factor may not have a strong influence in *Timema* as unresolved antagonisms are expected to be transient (54) and the stick insect X chromosome is very old and highly conserved (Fig. 1) and shares content with the X chromosome in other insect orders that diverged over 450 million years ago (33). Any maladaptive gene expression in females as a result of sexual conflict over dosage compensation would therefore be expected to be very small.

While our results show that dosage compensation is maintained in parthenogenetic species because selection in females against dosage compensation components is weak or because changes to the underlying mechanisms are constrained, we cannot formally exclude that perhaps not enough time has passed since the evolution of parthenogenesis to observe any changes to dosage compensation. Particularly under a fully neutral-drift scenario, timescales to observe decay could be longer for complex regulatory systems such as DC and MSCI (55, 56). Similarly, under a selective scenario (where there is selection against male-essential components of dosage compensation in females) a lack of time may be a plausible explanation in the young lineage *T. douglasi* and for the lack of apparent changes in dosage compensation in whole-body samples of males of parthenogenetic aphid lineages (57), however, it is very unlikely for the oldest species used here (*T. genevievae*) with an age estimate of over 1.5 million years (41, 42, 55).

### A stronger signature of MSCI in parthenogenetic than sexual species driven by autosomal mis-expression during meiosis

We have shown previously that the X chromosome is consistently inactivated throughout male meiosis in sexual *Timema* species (17, 44). Accordingly, we observe very low expression of X-linked genes in the testes, much lower than the expectation for a lack of dosage compensation (Fig. 3). We do however still observe some expression of the X in males, as expected given the presence of non-meiotic cells in testes. Surprisingly, we see a consistently lower expression of X-linked genes in males from parthenogenetic than sexual species (Fig. 3, S12). This could indicate variable cellular composition of testes such as a higher proportion of meiotic cells in parthenogenetic than sexual males, or, more likely, that autosomal expression is more persistent across meiosis in parthenogenetic as compared to sexual males. In sexual *Timema* males, there are two bursts of autosomal transcription during meiosis (44): a first at the end of prophase I (specifically, during pachytene and diplotene stages), and a second during meiosis II at the spermatid stage.

**Fig. 3:**
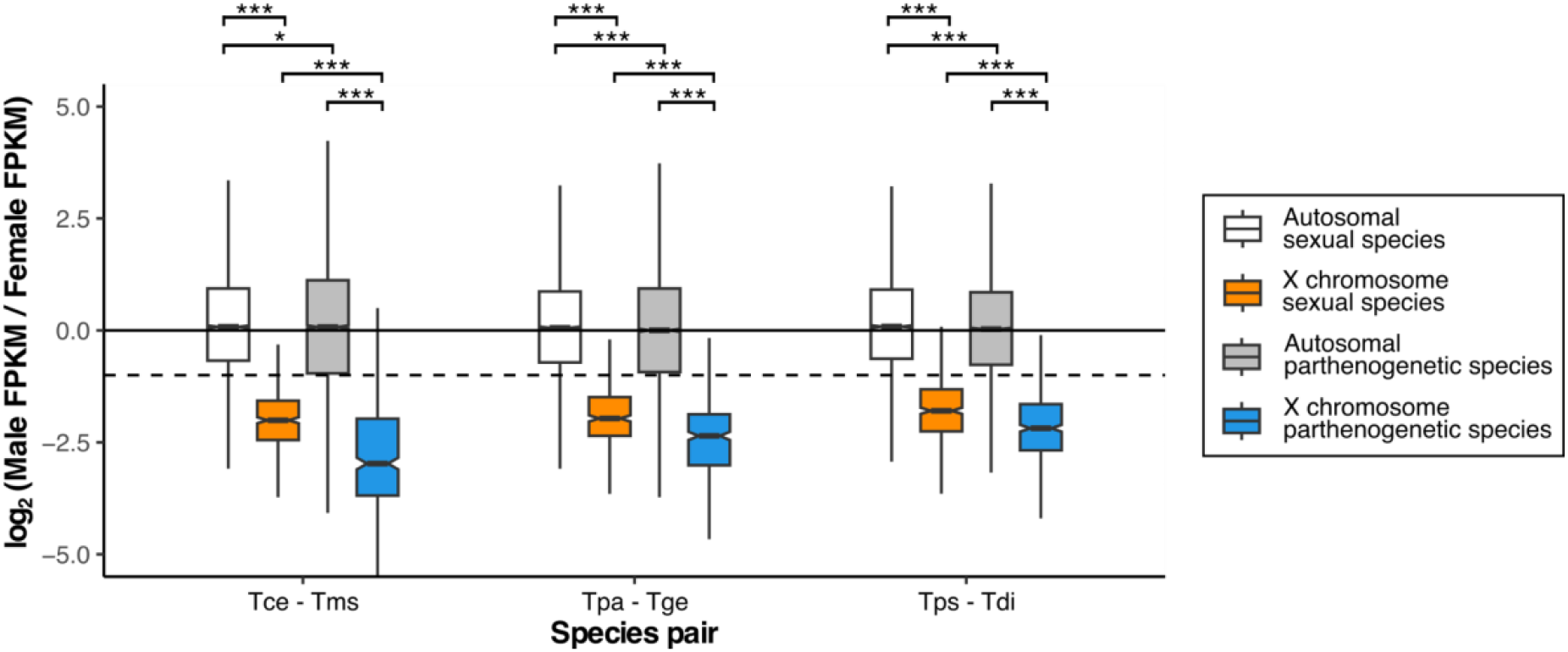
A stronger signature of MSCI in parthenogenetic than sexual species. Gene expression in reproductive tissues: Log_2_ of male to female expression ratio for the X and autosomes in testes and female gonads. Species codes: Tce = *T. cristinae*, Tms = *T. monikense*, Tpa = *T. podura*, Tge = *T. genevievae*, Tps = *T. poppense*, Tdi = *T. douglasi*. Asterisks indicate the significance (FDR) of Wilcoxon tests (***<0.001, **<0.01, *<0.05, NS > 0.05). Dashed lines represent a two-fold reduction in expression in males (as expected if there was no dosage compensation).

To detect potential changes to these temporal patterns of autosomal expression in parthenogenetic males, we performed immunolabeling of testis tissue of *T. monikense* and *T. douglasi* using RNA polymerase II, a conserved transcriptional marker. This labeling revealed that autosomal expression during prophase I is longer in parthenogenetic than sexual males. In the former, the RNA polymerase II signal already appears at the onset of prophase I (i.e., during the zygotene stage; Fig. 4, S13) and is maintained until diplotene (Fig. 4, S13), while in the latter it only appears at pachytene (44). Since males of parthenogenetic species are not exposed to selection, this extension of the autosomal transcription burst during prophase I is most likely a consequence of drift, i.e., it reflects autosomal mis-expression during meiosis in parthenogenetic males.

**Fig. 4:**
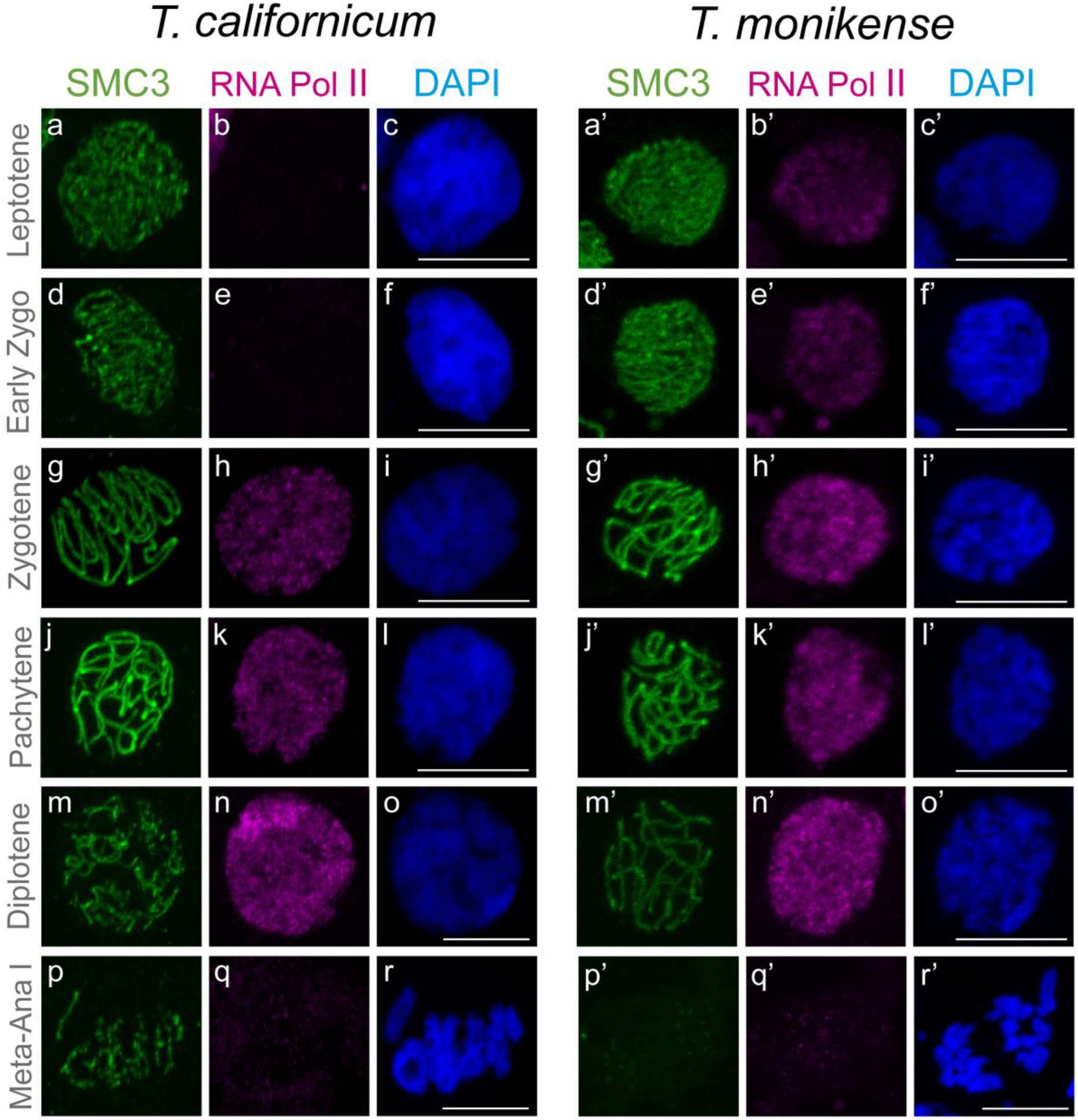
Misregulation of autosomal gene expression in meiotic cells of rare males produced by parthenogenetic females. RNApol-II distribution in meiocytes of sexual (*T. californicum*) and parthenogenetically produced (*T. monikensis*) males. Projections of stack images acquired across squashed meiocytes at the stages indicated, stained with DAPI (blue) and double immunolabeled for SMC3 (green) and RNApol-II (magenta). RNApol-II labelling is already present at leptotene and early zygotene in parthenogenetic *T. monikensis* males, while it only appears during late zygotene in sexual *T. californicum* males, as shown previously (Dumas et al 2025). Note that the used immunolabeling allows for inference of qualitative (presence-absence) rather than quantitative (intensity of signals) differences (Fig. S13). Scale bar: 10 μm. Corresponding image analyses for males of *T. douglasi* are available in Fig. S13).

## Conclusion

Our results demonstrate that both dosage compensation and meiotic sex chromosome inactivation (MSCI) are robustly maintained in parthenogenetic stick insects, despite the long-term absence of selective pressures typically associated with male phenotypes. This maintenance suggests that the molecular mechanisms underlying dosage compensation and MSCI are either functionally constrained (e.g. due to pleiotropic effects in females), maintained by weak purifying selection (for example through sufficiently frequent reproduction of rare males), or decay too slowly as a consequence of drift for phenotypic effects to be observed in the stick insects. Nevertheless, the enhanced signature of MSCI in parthenogenetic males appears to be driven by extended autosomal transcription during meiosis, revealing an unexpected consequence of relaxed selection on gene expression. Together, these findings highlight the stability and evolutionary inertia of complex regulatory processes, even in the face of dramatic shifts in reproductive mode.

## Methods

### Reference genome assemblies

Reference genome assemblies were produced for *T. cristinae* (Bioproject PRJNA1268975/ Accession JBOIUC000000000) and *T. podura* (Bioproject PRJNA1268975/ Accession JBOIUD000000000) from wild collected females (GPS coordinates 34°30’53.2”N, 119°46’45.3”W and 33°47’47.0”N, 116°46’34.6”W). Contigs for the *T. cristinae* assembly were based on PacBio sequencing (43x) and on a combination of Nanopore and Illumina sequencing for *T. podura* (sequenced at 85x respectively 62x coverage). Scaffolding was done using a Hi-C library (sequenced at approximately 69 x and 53 x coverage for *T. cristinae* and *T. podura* respectively) to produce the final reference assembly.

#### DNA extraction and library preparation

To extract high molecular weight (HMW) DNA, we flash-froze a single female (without gut) in liquid nitrogen and ground it using a Cryomill (Retsch). We then extracted HMW DNA using a G/20 Genomic Tips kit (Qiagen) following the manufacturer’s protocols and checked DNA integrity on a pulse field agarose gel.

For *T. cristinae*, a PacBio library was prepared using the Template Prep Kit 1.0 following manufacturer’s instructions and sequenced on a single cell of the PacBio SMRT™ system. For *T. podura*, four ONT libraries were prepared following Oxford Nanopore instructions. A first library was prepared using the SQK-LSK108 ligation sequencing kit and was loaded on a MinION R9.4 Flow Cells, and then three libraries were prepared using the SQK-LSK109 ligation sequencing kit and were loaded on three PromethION R9.4.1 Flow Cells. A PCR free Illumina library (based on the same HMW extraction) was prepared using the Kapa Hyper Prep Kit (Roche, Basel, Switzerland), following manufacturer instructions. The library was quantified by qPCR using the KAPA Library Quantification Kit for Illumina Libraries (Roche), and the library profile was assessed using a High Sensitivity DNA kit on an Agilent Bioanalyzer (Agilent Technologies, Santa Clara, CA, USA). The library was then sequenced on an Illumina HiSeq 4000 instrument (Illumina, San Diego, CA, USA), using 150 base-length read chemistry in a paired-end mode.

A Hi-C library was prepared for each species using the Proximo Hi-C Kit. We generated ground tissue for cross-linking using Cryomill (Retsch) and following manufacturer instructions, using a different female than the one used for Nanopore and HiSeq sequencing, from the same natural population. Library construction and sequencing (250 million read pairs) was outsourced to Phase Genomics (Seattle).

#### Assembly pipelines

We used different strategies for genome assembly for the two species. For *T. cristinae* raw PacBio reads were assembled into contigs using Canu (58) with the parameter correctedErrorRate=0.105. For *T. podura* raw Oxford Nanopore reads were filtered using Filtlong v0.2.0 (https://github.com/rrwick/Filtlong) with the parameters --min_length 1000 -- keep_percent 90 --target_bases 69050000000. The filtered Nanopore reads were then assembled into contigs using Flye v2.8.1 (59) with --genome-size 1.381g. All Nanopore reads were mapped against the contigs using minimap2 v2.19 (60) with the parameters -c -x map-ont and a first step of polishing was performed using Racon v1.4.3 (61). Three additional rounds of polishing were conducted: Illumina short reads were aligned to the contigs using BWA mem v0.7.17 (62) and polishing was performed using Pilon v1.23 (63).

Both assemblies were then decontaminated using BlobTools v1.0 (64) under the taxrule “bestsumorder”. Hit files were generated after a blastn v2.10.1+ against the NBCI nt database, searching for hits with an e-value below 1e-25 (parameters: -max_target_seqs 10 -max_hsps 1 -evalue 1e-25). Contigs without hits to metazoans were removed from the assemblies. Haplotypic duplications were then filtered out as follows. Filtered reads were then mapped against the decontaminated genomes using minimap2 and haplotigs were detected with Purge Haplotigs v1.1.1 (65) using the parameters -l 3 -m 27 -h 190 -j 101 --editor-coarse-resolution 250000 for *T. cristinae* and -l 3 -m 45 -h 195 -j 101 --editor-coarse-resolution 50000 for *T. podura*.

Hi-C reads from each species were then mapped against their respective haploid genome using Juicer v1.6 (66) with the restriction site Sau3AI. Finally, chromosome-level scaffolding was performed using 3D-DNA v180922 (67) with species-specific parameters, as recommended by the authors. The resulting Hi-C contact matrices were visualized with Juicebox, and minor polishing was conducted in accordance with these recommendations. The completeness of both genome assemblies was assessed with BUSCO v5.1.2 (68) against the *insecta_odb10* dataset using the --long and --augustus parameters.

#### Annotation

We annotated the *T. cristinae* and *T. podura* genomes in a similar way as we previously annotated the *T. poppense* genome (17), using a combination of *ab initio* gene prediction, protein homology, and RNA-seq using the Braker2 pipeline (v. 2.1.6) (69). Firstly, each genome assembly was soft-masked using RepeatModeler (v. 2.0.2, options: -LTRStruct, - engine ncbi) (70) and RepeatMasker (v. 4.1.2, options: -engine ncbi, -xsmall) (71). For protein evidence, we used the arthropod protein sequences from OrthoDB (v.10.1) (72) and the predicted protein sequences for *Timema* from our previous genome assemblies. For RNAseq evidence, we used the RNAseq libraries generated here for *T. cristinae* and *T. monikense* along with publicly available data (163 libraries generated here, 43 public (Bioproject accessions: PRJNA380865, PRJNA679785, PRJNA1295360, PRJNA504764 (42, 43, 73))) to annotate the *T. cristinae* genome and for *T. podura* and *T. genevievae* (119 libraries generated here, 29 public (Bioproject accessions: PRJNA380865, PRJNA679785, PRJNA1295360 (42, 43))) to annotate the *T. podura* genome. This is a total of 354 RNAseq libraries (330 paired-end, 24 single-end) covering over 19 different life stages, tissue, and sex combinations for each species (see Table S3). Reads were quality trimmed with Trimmomatic (v. 0.39, options: ILLUMINACLIP:3:25:6 LEADING:9 TRAILING:9 SLIDINGWINDOW:4:15 MINLEN:80) (74) before mapping to the genome assembly with STAR (v. 2.7.8a, options: --twopassMode Basic) (75). Braker2 was run using protein evidence and RNAseq separately with the gene predictors Augustus (v. 3.4.0) (76) and Genemark (v. 4.72) (77). Following the RNAseq run, UTR predictions were added to the RNAseq gene predictions using GUSHR (v. 1.0) in Braker2 (--addUTR=on). The separate gene predictions were then merged using TSEBRA (v1.0.3) using the pref_braker1.cfg configuration file, which weights RNAseq evidence more strongly than the default option. We then ran BUSCO (v. 5.3.2, insecta_odb10) on the gene regions annotated by Braker2 and on the whole genome assembly. Any genes found by BUSCO but missed by Braker were then added to the annotation (69 genes for *T. cristinae* and 19 genes for *T. podura*). ncRNA genes were predicted using Infernal (v. 1.1.2, minimum e-value 1e-10) (78). GO-terms for protein-coding gene predictions were obtained using blastP within OmicsBox (v3.1.2, default parameters) to blast the nr *Drosophila melanogaster* database (taxonomy filter: 7227). 1-to-1 orthologs between were obtained for protein-coding genes using OrthoFinder (version 2.5.5) (79) with default options.

#### Identification of the X chromosome

We used a coverage approach to identify X-linked scaffolds in our new genome assemblies. To do this we mapped publicly available Illumina genome reads from males and females (4 males and 5 females for *T. cristinae* and 4 males and 2 females for *T. podura* (PRJNA670663, PRJNA725673 (11, 43)) to the respective genome assemblies and their publicly available mtDNA genomes (Accessions) using BWA-MEM (v. 0.7.15). Multi-mapping and poor quality alignments were filtered (removing reads with XA:Z or SA:Z tags or a mapq <30). PCR duplicates were removed with Picard (v. 2.9.0) (http://broadinstitute.github.io/picard/). Coverage was then estimated for 100000 bp sliding windows using BEDTools (v. 2.26.0) (80). To compare coverage between males and females, coverage was first summed for all male and all female libraries per window. Male and female coverage was then normalized by modal coverage to adjust for differences in overall coverage. X-linked windows were then identified using the log_2_ ratio of male to female coverage. Autosomal windows should have equal coverage in males and females (log2 ratio of male to female coverage ≈ 0) and X-linked windows should have half the coverage in males as in females (log2 ratio of male to female coverage ≈ −1). This approach identified one large scaffold in each assembly as the X chromosome (Fig. S14-16).

We then repeated these analyses using genome reads from each of the parthenogenetic species. Female reads were publicly available (5 for *T. monikense*, 6 for *T. genevievae*, and 5 for *T. douglasi*, PRJNA670663 (43)) as were the two male samples for *T. douglasi* (PRJNA808673, (81)). For *T. monikense* and *T. genevievae* we also generated new Illumina libraries for males (2 for *T. genevievae* and 4 for *T. monikense*) and an additional female library for *T. genevievae* (Accession number: PRJNA1293928) as described previously (43) using the carcasses after tissue dissection described below. Reads were then mapped to the genome of each sexual sister species as described above (the sexual species genome for *T. douglasi* is our previously published genome *T. poppense*, Accession number: PRJNA1126215 (17)). Each of the newly generated samples were checked for normal coverage distributions before calculating coverage ratios.

#### RNA extraction and sequencing

We generated a total of 282 RNA-seq libraries for eight female tissues (antennae, brain, defence glands, fat body, femur, tarsi, and gonads) and nine male tissues (the same as the female plus the male accessory glands) for *T. cristinae, T. podura, T. monikense*, and *T. genevievae* following our previously developed methodology (17) (see Table S3 for details). Individuals were collected as nymphs in the field and reared to adulthood in the laboratory. Prior to dissection all individuals were isolated in Petri dishes and fed on an artificial diet (a mix of flour, sugar, and water) for two days, allowing their gut contents to be cleared of plant material. Insects were then anesthetized with CO_2_, dissected, and tissues individually flash-frozen in liquid nitrogen before being stored at -80°C.

TRIzol solution (1 mL) and a small amount of ceramic beads (Sigmund Lindner) were added to each tissue. Samples were homogenized using a tissue homogenizer (Precellys Evolution; Bertin Technologies). Chloroform (200 μL) was added to each sample and samples were then vortexed for 15 seconds and centrifuged for 25 minutes at 12,000 rpm at 4°C. The upper phase containing the RNA was then transferred to a new 1.5 mL tube with the addition of isopropanol (650 μL). The samples were vortexed and placed at -20°C overnight. Samples were then centrifuged for 30 minutes at 12,000 rpm at 4°C. The liquid supernatants were removed, and the RNA pellet underwent two washes using 80% and 70% ethanol. Each wash was followed by a 5-minute centrifugation step at 12,000 rpm. Finally, the RNA pellet was resuspended in nuclease-free water and quantified using a fluorescent RNA-binding dye (QuantiFluor RNA System) and nanodrop (DS-11 FX).

Library preparation using NEBNext (New England BioLabs) and sequencing on an Illumina NovaSeq 6000 platform with 100 bp paired-end sequencing (36.2 million read pairs per sample on average) was outsourced to a sequencing facility (Fasteris, Geneva).

#### Gene expression analyses

We mapped RNAseq reads from *T. cristinae, T. podura, T. monikense*, and *T. genevievae* generated here and reads for *T. poppense* and *T. douglasi* we generated previously (17) to their respective sexual species genomes as described above. This gave 4 replicates for each of the 16 sex and tissue combinations for each of the six species, except for the male samples of parthenogenetic species (*T. monikense, T. genevievae*, and *T. douglasi*) where we had fewer replicates due to the difficulty of obtaining these specimens (n = 2-4 see Table S4 for details). Read counts were obtained using HTSeq v.0.9.1 (82), with the following options (htseq-count --order = name --type = exon --idattr = gene_id --stranded = reverse). Expression analyses were performed using the Bioconductor package EdgeR (v 3.42.4) (83), and done separately for each species and tissue. Normalization factors for each library were computed using the TMM method and were used to calculate normalized expression levels (FPKM, Fragments Per Kilobase of transcript per Million mapped reads). mtDNA, rRNA, tRNA and genes with low expression (less than 2 FPKM in 2 or more libraries per sex in either the sexual or parthenogenetic sister species) were excluded. This filtering was done to ensure that genes used for gene expression comparisons between sexual and parthenogenetic sister species and between males and females were the same. To examine dosage compensation, we used the log_2_ ratio of mean male expression level to female expression level and used Wilcoxon tests (adjusted for multiple testing using Benjamini and Hochberg’s (84) algorithm) to determine if differences in expression between the autosomes and the X chromosome were significant.

#### Immunolabeling of parthenogenetic male meiotic cells

In order to determine the dynamics of autosomal gene expression during meiosis in parthenogenetic males, we followed the same protocol previously used to characterize autosomal gene expression during meiosis in sexual males (44). In brief, we performed double immunolabeling of SMC3_488 (#AB201542, AbCam) as a marker of cohesin axes (this allows for the assessment of synapsis progression, and hence, cell cycle stage) and RNA polymerase II phosphorylated at serine 2 (p-RNApol-II (ab193468, AbCam); an indicator of transcription). Gonads from adult *T. monikense* and *T. douglasi* males were dissected in 1X PBS, then fixed in paraformaldehyde (2%) and Triton X-100 (0.1%) solution for 15 minutes. Gonads were then squashed on slides coated with poly-L-lysine, and flash frozen in liquid nitrogen. Slides were incubated in PBS 1x for 20 minutes at room temperature, then blocked with a BSA 3% solution (prepared by dissolving 3g of BSA in 100mL of PBS 1X) for 20 minutes. Slides were then incubated overnight at 4°C in a humid chamber with the primary (p-RNA-Pol II), diluted in BSA 3% (1:100), and washed 3x for 5 minutes in PBS 1X. The secondary antibody, anti-rabbit Alexa 594 (711-585-152, Jackson), diluted in BSA 3% (1:100), was applied, and samples were incubated for 45 minutes at room temperature. Further washes were conducted as described above, followed by an extended 10-minute wash in PBS 1X, and by blocking using a 5% Normal Rabbit Serum (NRS) solution (X090210-8, Agilent Technology) for 30 minutes at room temperature. Another 5-minute wash in PBS 1x was performed to remove excess blocking solution. Samples were then incubated with the labelled antibody SMC3_488 (#AB201542, AbCam) at a 1:100 dilution for 1 hour at room temperature, followed by a final series of washes and staining with DAPI (D9542, Sigma-Aldrich) for 3 minutes at room temperature. Finally, after a 5-minute wash in 1x PBS, two drops of Vectashield were added to the slide, which was then covered with a coverslip, and sealed with nail varnish for imaging. Image acquisition was performed at the Cellular Imaging Facility (CIF, University of Lausanne) using a Zeiss LSM 880 Airyscan.

## Data

Genome assemblies have been deposited at DDBJ/ENA/GenBank under the accessions JBOIUC000000000 (*T. cristinae*) and JBOIUD000000000 (*T. podura*). The Illumina and PromethION sequencing data for *T. podura* are available in the European Nucleotide Archive under the following project PRJEB94285. Male genomic reads have been deposited at SRA under the following project PRJNA1293928. RNAseq reads have been deposited at SRA under the following projects: PRJNA1320679, PRJNA1327219. Code and annotations are available here: https://github.com/DarrenJParker/Timema_DC_MSCI_code and will be archived upon acceptance.

## Acknowledgments

This work was supported by the European Research Council Consolidator Grant 864672 (“No Sex No Conflict”, T.S.), Swiss FNS grant 31003A_182495 (T.S.), the Genoscope, the Commissariat à l’Énergie Atomique et aux Énergies Alternatives (CEA) and France Génomique (ANR-10-INBS-09–08). We thank the Lausanne Genomic Technologies Facility for PacBio library preparation and sequencing.

## Notes

### Competing Interest Statement

The authors have declared no competing interest.

### Summary of Updates

updated clarity, additional analyses and cell bio

## References

1. D. Bachtrog, et al., Sex determination: why so many ways of doing it? PLoS Biol. 12, e1001899 (2014).

2. C. M. Disteche, Dosage compensation of the sex chromosomes. Annu. Rev. Genet. 46, 537–560 (2012).

3. L. Gu, J. R. Walters, Evolution of sex chromosome dosage compensation in animals: a beautiful theory, undermined by facts and bedeviled by details. Genome Biol. Evol. 9, 2461–2476 (2017).

4. J. E. Mank, Sex chromosome dosage compensation: definitely not for everyone. Trends Genet. 29, 677–683 (2013).

5. N. Brockdorff, B. M. Turner, Dosage compensation in mammals. Cold Spring Harb. Perspect. Biol. 7, a019406 (2015).

6. R. Marin, et al., Convergent origination of a Drosophila-like dosage compensation mechanism in a reptile lineage. Genome Res. 27, 1974–1987 (2017).

7. D. C. H. Metzger, B. A. Sandkam, I. Darolti, J. E. Mank, Rapid Evolution of Complete Dosage Compensation in Poecilia. Genome Biol. Evol. 13 (2021).

8. I. Darolti, et al., Extreme heterogeneity in sex chromosome differentiation and dosage compensation in livebearers. Proc. Natl. Acad. Sci. U. S. A. 116, 19031–19036 (2019).

9. B. J. Meyer, Sex in the worm: counting and compensating X-chromosome dose. Trends Genet. 16, 247–253 (2000).

10. P. Chauhan, et al., Genome assembly, sex-biased gene expression and dosage compensation in the damselfly Ischnura elegans. Genomics 113, 1828–1837 (2021).

11. D. J. Parker, K. S. Jaron, Z. Dumas, M. Robinson-Rechavi, T. Schwander, X chromosomes show relaxed selection and complete somatic dosage compensation across Timema stick insect species. J. Evol. Biol. (2022). 10.1111/jeb.14075.

12. A. Pal, B. Vicoso, The X Chromosome of Hemipteran insects: conservation, dosage compensation and sex-biased expression. Genome Biol. Evol. 7, 3259–3268 (2015).

13. J. G. Rayner, T. J. Hitchcock, N. W. Bailey, Variable dosage compensation is associated with female consequences of an X-linked, male-beneficial mutation. Proc. Biol. Sci. 288, 20210355 (2021).

14. E. G. Prince, D. Kirkland, J. P. Demuth, Hyperexpression of the X chromosome in both sexes results in extensive female bias of X-linked genes in the flour beetle. Genome Biol. Evol. 2, 336–346 (2010).

15. B. Vicoso, D. Bachtrog, Numerous transitions of sex chromosomes in Diptera. PLoS Biol. 13, e1002078 (2015).

16. T. Conrad, A. Akhtar, Dosage compensation in Drosophila melanogaster: epigenetic fine-tuning of chromosome-wide transcription. Nat. Rev. Genet. 13, 123–134 (2012).

17. J. Djordjevic, et al., Dynamics of X chromosome hyper-expression and inactivation in male tissues during stick insect development. PLoS Genet. 21, e1011615 (2025).

18. J. E. Mank, H. Ellegren, All dosage compensation is local: gene-by-gene regulation of sex-biased expression on the chicken Z chromosome. Heredity 102, 312–320 (2009).

19. P. S. Burgoyne, S. K. Mahadevaiah, J. M. A. Turner, The consequences of asynapsis for mammalian meiosis. Nat. Rev. Genet. 10, 207–216 (2009).

20. B. D. McKee, M. A. Handel, Sex chromosomes, recombination, and chromatin conformation. Chromosoma 102, 71–80 (1993).

21. A. I. Kalita, C. I. Keller Valsecchi, Dosage compensation in non-model insects-progress and perspectives. Trends Genet. (2024). 10.1016/j.tig.2024.08.010.

22. J. B. Pease, M. W. Hahn, Sex chromosomes evolved from independent ancestral linkage groups in winged insects. Mol. Biol. Evol. 29, 1645–1653 (2012).

23. F. N. Hamada, P. J. Park, P. R. Gordadze, M. I. Kuroda, Global regulation of X chromosomal genes by the MSL complex in Drosophila melanogaster. Genes Dev. 19, 2289–2294 (2005).

24. T. Straub, G. D. Gilfillan, V. K. Maier, P. B. Becker, The Drosophila MSL complex activates the transcription of target genes. Genes Dev. 19, 2284–2288 (2005).

25. A. I. Kalita, et al., The sex-specific factor SOA controls dosage compensation in Anopheles mosquitoes. Nature 623, 175–182 (2023).

26. E. Krzywinska, P. Ribeca, L. Ferretti, A. Hammond, J. Krzywinski, A novel factor modulating X chromosome dosage compensation in Anopheles. Curr. Biol. 33, 4697–4703.e4 (2023).

27. R. J. Davis, E. J. Belikoff, E. H. Scholl, F. Li, M. J. Scott, No blokes is essential for male viability and X chromosome gene expression in the Australian sheep blowfly. Curr. Biol. 28, 1987–1992.e3 (2018).

28. M. A. Toups, B. Vicoso, Insect sex chromosome evolution: conservation, turnover, and mechanisms of dosage compensation. Curr. Opin. Insect Sci. 101411 (2025).

29. N. Page, et al., Single-cell profiling of Anopheles gambiae spermatogenesis defines the onset of meiotic silencing and premeiotic overexpression of the X chromosome. Commun. Biol. 6, 850 (2023).

30. S. Schoenmakers, et al., Female meiotic sex chromosome inactivation in chicken. PLoS Genet. 5, e1000466 (2009).

31. S. H. Namekawa, J. T. Lee, XY and ZW: is meiotic sex chromosome inactivation the rule in evolution? PLoS Genet. 5, e1000493 (2009).

32. R. L. Kelley, et al., Expression of msl-2 causes assembly of dosage compensation regulators on the X chromosomes and female lethality in Drosophila. Cell 81, 867–877 (1995).

33. M. A. Toups, B. Vicoso, The X chromosome of insects likely predates the origin of class Insecta. Evolution 77, 2504–2511 (2023).

34. J. T. Lee, Disruption of imprinted X inactivation by parent-of-origin effects at Tsix. Cell 103, 17–27 (2000).

35. C. J. van der Kooi, T. Schwander, On the fate of sexual traits under asexuality. Biol. Rev. Camb. Philos. Soc. 89, 805–819 (2014).

36. J. M. Belote, J. C. Lucchesi, Control of X chromosome transcription by the maleless gene in Drosophila. Nature 285, 573–575 (1980).

37. E. Lifschytz, D. L. Lindsley, The role of X-chromosome inactivation during spermatogenesis (Drosophila-allocycly-chromosome evolution-male sterility-dosage compensation). Proc. Natl. Acad. Sci. U. S. A. 69, 182–186 (1972).

38. Z. Bao, A. Belyi, E. Argyridou, J. Parsch, Disruption of meiotic sex chromosome inactivation by X-autosome translocations in Drosophila melanogaster. Genetics 232, iyag011.(2026).

39. C. C. Avelino, et al., Meiotic Sex Chromosome Inactivation: Conservation across the Drosophila genus. PLoS Genet. 21, e1011511 (2025).

40. T. Schwander, B. J. Crespi, Multiple direct transitions from sexual reproduction to apomictic parthenogenesis in Timema stick insects. Evolution 63, 84–103 (2009).

41. T. Schwander, L. Henry, B. J. Crespi, Molecular evidence for ancient asexuality in Timema stick insects. Curr. Biol. 21, 1129–1134 (2011).

42. J. Bast, et al., Consequences of asexuality in natural populations: insights from stick insects. Mol. Biol. Evol. 35, 1668–1677 (2018).

43. K. S. Jaron, et al., Convergent consequences of parthenogenesis on stick insect genomes. Science Advances 8, eabg3842 (2022).

44. Z. Dumas, W. Toubiana, M. Delattre, T. Schwander, Dynamics of recombination, X inactivation and centromere proteins during stick insect spermatogenesis. bioRxiv 2024.09.05.611401 (2024).

45. T. Schwander, B. J. Crespi, R. Gries, G. Gries, Neutral and selection-driven decay of sexual traits in asexual stick insects. Proc. Biol. Sci. 280, 20130823 (2013).

46. S. Freitas, D. J. Parker, M. Labédan, Z. Dumas, T. Schwander, Evidence for cryptic sex in parthenogenetic stick insects of the genus Timema. Proceedings of the Royal Society B: Biological Sciences 290, 20230404 (2023).

47. S. Simon, et al., Old World and New World Phasmatodea: phylogenomics resolve the evolutionary history of stick and leaf Insects. Frontiers in Ecology and Evolution 7, 345 (2019).

48. S. Bank, S. Bradler, A second view on the evolution of flight in stick and leaf insects (Phasmatodea). BMC Ecol. Evol. 22, 62 (2022).

49. X. Li, J. E. Mank, L. Ban, The grasshopper genome reveals long-term gene content conservation of the X Chromosome and temporal variation in X Chromosome evolution. Genome Res. 34, 997–1007 (2024).

50. Q.-L. Hu, et al., Chromosome-level assembly, dosage compensation and sex-biased gene expression in the small brown planthopper, Laodelphax striatellus. Genome Biol. Evol. 14, evac160 (2022).

51. A. M. Rice, Y. Li, P. Donnelly, A. McLysaght, Evolution of dosage-sensitive genes by tissue-restricted expression changes. Genome Biol. Evol. 17, evaf132 (2025).

52. J. Larsson, M. J. Svensson, P. Stenberg, M. Mäkitalo, Painting of fourth in genus Drosophila suggests autosome-specific gene regulation. Proc. Natl. Acad. Sci. U. S. A. 101, 9728–9733 (2004).

53. J. E. Mank, D. J. Hosken, N. Wedell, Some inconvenient truths about sex chromosome dosage compensation and the potential role of sexual conflict. Evolution 65, 2133–2144 (2011).

54. E. Flintham, V. Savolainen, S. P. Otto, M. Reuter, C. Mullon, The maintenance of genetic polymorphism underlying sexually antagonistic traits. Evol. Lett. 9, 259–272 (2025).

55. D. C. Lahti, et al., Relaxed selection in the wild. Trends Ecol. Evol. 24, 487–496 (2009).

56. D. J. Futuyma, Evolutionary constraint and ecological consequences. Evolution 64, 1865–1884 (2010).

57. H. Defendini, et al., The release of sexual conflict after sex loss is associated with evolutionary changes in gene expression. Proc. Biol. Sci. 292, 20242631 (2025).

58. S. Koren, et al., Canu: scalable and accurate long-read assembly via adaptive k-mer weighting and repeat separation. Genome Res. 27, 722–736 (2017).

59. M. Kolmogorov, J. Yuan, Y. Lin, P. A. Pevzner, Assembly of long, error-prone reads using repeat graphs. Nat. Biotechnol. 37, 540–546 (2019).

60. H. Li, Minimap2: pairwise alignment for nucleotide sequences. Bioinformatics 34, 3094–3100 (2018).

61. R. Vaser, I. Sović, N. Nagarajan, M. Šikić, Fast and accurate de novo genome assembly from long uncorrected reads. Genome Res. 27, 737–746 (2017).

62. H. Li, Aligning sequence reads, clone sequences and assembly contigs with BWA-MEM. arXiv 1303.3997 (2013).

63. B. J. Walker, et al., Pilon: an integrated tool for comprehensive microbial variant detection and genome assembly improvement. PLoS One 9, e112963 (2014).

64. D. R. Laetsch, M. L. Blaxter, BlobTools: Interrogation of genome assemblies. F1000Res. 6, 1287 (2017).

65. M. J. Roach, S. A. Schmidt, A. R. Borneman, Purge Haplotigs: allelic contig reassignment for third-gen diploid genome assemblies. BMC Bioinformatics 19, 460 (2018).

66. N. C. Durand, et al., Juicer provides a one-click system for analyzing loop-resolution Hi-C experiments. Cell Syst. 3, 95–98 (2016).

67. O. Dudchenko, et al., De novo assembly of the Aedes aegypti genome using Hi-C yields chromosome-length scaffolds. Science 356, 92–95 (2017).

68. M. Seppey, M. Manni, E. M. Zdobnov, BUSCO: Assessing genome assembly and annotation completeness. Methods Mol. Biol. 1962, 227–245 (2019).

69. T. Brůna, K. J. Hoff, A. Lomsadze, M. Stanke, M. Borodovsky, BRAKER2: automatic eukaryotic genome annotation with GeneMark-EP+ and AUGUSTUS supported by a protein database. NAR Genom Bioinform 3, lqaa108 (2021).

70. J. M. Flynn, et al., RepeatModeler2 for automated genomic discovery of transposable element families. Proc. Natl. Acad. Sci. U. S. A. 117, 9451–9457 (2020).

71. A. Smith, R. Hubley, P. Green, RepeatMasker Open-4.0. RepeatMasker Open-4.0 (2013).

72. E. V. Kriventseva, et al., OrthoDB v10: sampling the diversity of animal, plant, fungal, protist, bacterial and viral genomes for evolutionary and functional annotations of orthologs. Nucleic Acids Res. 47, D807–D811 (2019).

73. D. J. Parker, J. Djordjevic, T. Schwander, Olfactory proteins in Timema stick insects. Frontiers in Ecology and Evolution 7, 101 (2019).

74. A. M. Bolger, M. Lohse, B. Usadel, Trimmomatic: a flexible trimmer for Illumina sequence data. Bioinformatics 30, 2114–2120 (2014).

75. A. Dobin, et al., STAR: ultrafast universal RNA-seq aligner. Bioinformatics 29, 15–21 (2013).

76. M. Stanke, et al., AUGUSTUS: ab initio prediction of alternative transcripts. Nucleic Acids Res. 34, W435–9 (2006).

77. T. Brůna, A. Lomsadze, M. Borodovsky, GeneMark-EP+: eukaryotic gene prediction with self-training in the space of genes and proteins. NAR Genom Bioinform 2, qaa026 (2020).

78. E. P. Nawrocki, S. R. Eddy, Infernal 1.1: 100-fold faster RNA homology searches. Bioinformatics 29, 2933–2935 (2013).

79. D. M. Emms, S. Kelly, OrthoFinder: phylogenetic orthology inference for comparative genomics. Genome Biol. 20, 238 (2019).

80. A. R. Quinlan, I. M. Hall, BEDTools: a flexible suite of utilities for comparing genomic features. Bioinformatics 26, 841–842 (2010).

81. C. Larose, G. Lavanchy, S. Freitas, D. J. Parker, T. Schwander, Facultative parthenogenesis in an asexual stick insect: re-evolution of sex or vestigial sexual capacity? bioRxiv 2022.03.25.485836(2022).

82. S. Anders, P. T. Pyl, W. Huber, HTSeq-a Python framework to work with high-throughput sequencing data. Bioinformatics 31, 166–169 (2015).

83. M. D. Robinson, D. J. McCarthy, G. K. Smyth, edgeR: a Bioconductor package for differential expression analysis of digital gene expression data. Bioinformatics 26, 139–140 (2010).

84. Y. Benjamini, Y. Hochberg, Controlling the false discovery rate: A practical and powerful approach to multiple testing. J. R. Stat. Soc. Series B Stat. Methodol. 57, 289–300 (1995).

