## Supplementary material for "Dosage compensation and meiotic sex chromosome inactivation are maintained in the absence of selection": Supp material

##### *Identifying the X chromosome*

The X chromosome was identified using a coverage approach. Since the X chromosome is present in two copies in females and one in males, X-linked scaffolds will have half the coverage in males when compared to females. Using this approach we identified a large scaffold in each of the genome assemblies as the X chromosome (Fig. S14-16). The size of the X chromosome was similar for each species (10-12% of the genome, Table S5) with a similar number of genes on the X for each species (Tables S17-19). Repeating these analyses using samples from the parthenogenetic sister species (mapped to the sexual genome) identified the same scaffold as the X chromosome, with a similar reduction of coverage in the males (Fig. S17-S19).

#### Supplementary figures

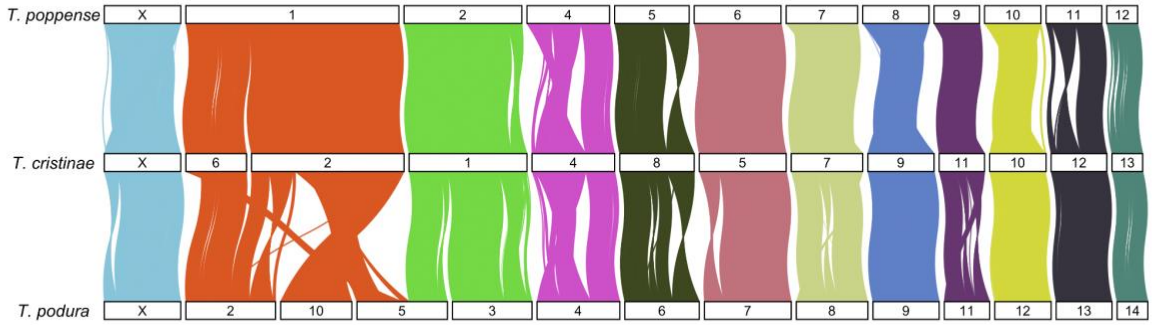


**Fig. S1 | Synteny between 8118 one-to-one orthologs.**

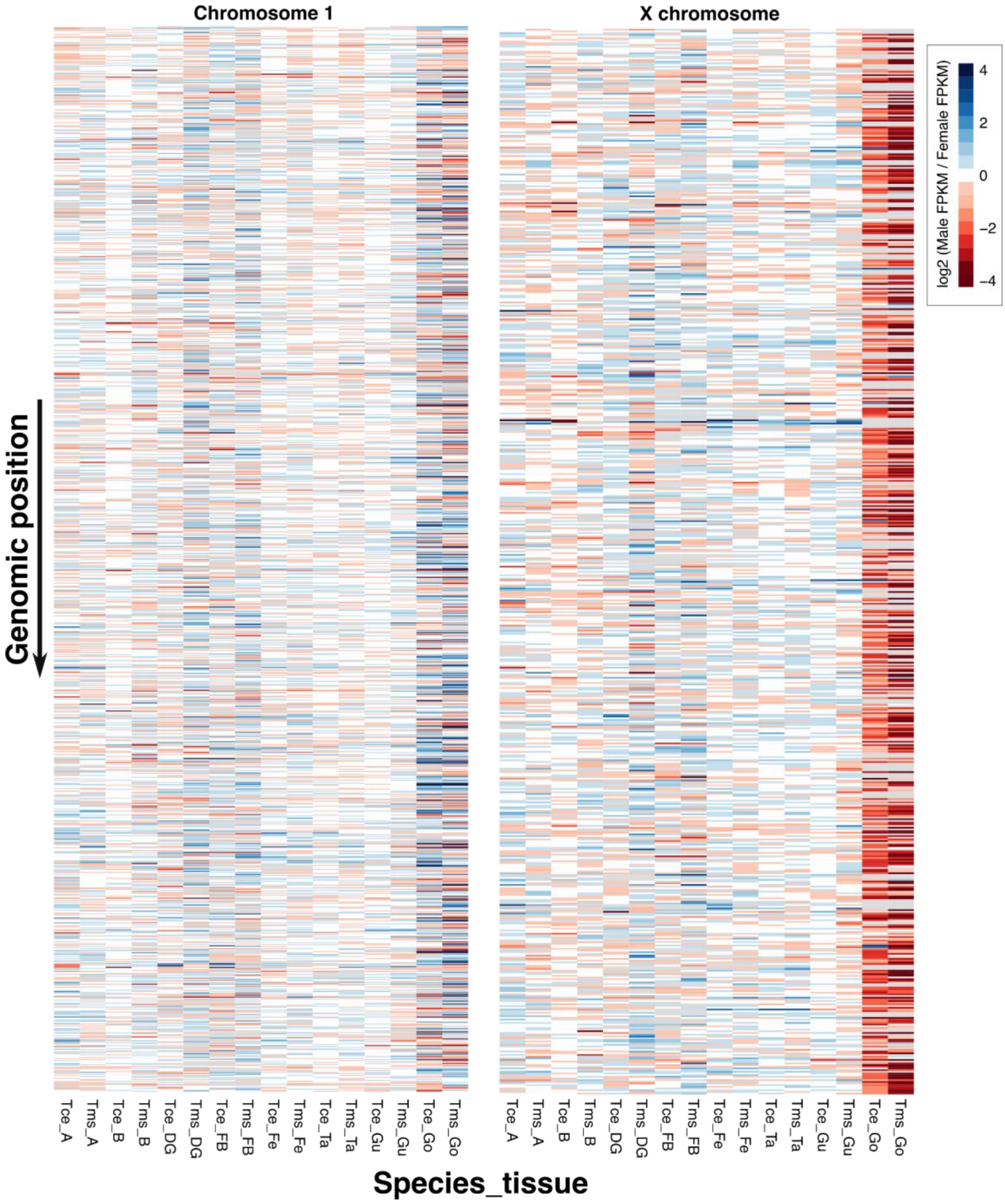


**Fig. S2 | Male to female expression ratios along the X and chromosome 1 for *T. cristinae* (Tce) and *T. monikense* (Tms) in Antennae (A), Brain (B), Defence glands (DG), Fat body (FB), Femur (Fe), Gut (Gu), Tarsi (Ta), and Gonads (Go).**

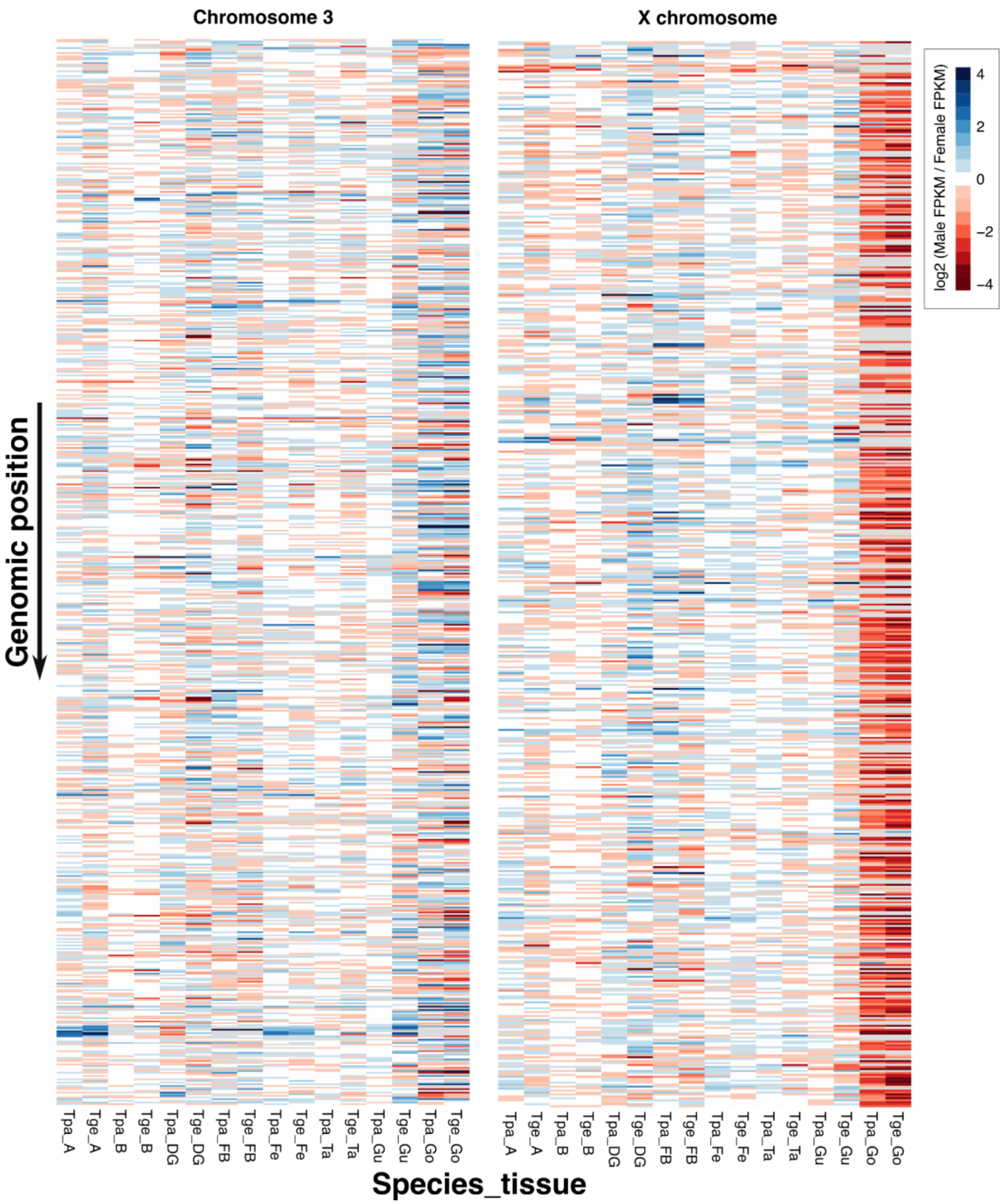


**Fig. S3 | Male to female expression ratios along the X and chromosome 3 for *T. podura* (Tpa) and *T. genevievae* (Tge) in Antennae (A), Brain (B), Defence glands (DG), Fat body (FB), Femur (Fe), Gut (Gu), Tarsi (Ta), and Gonads (Go).**

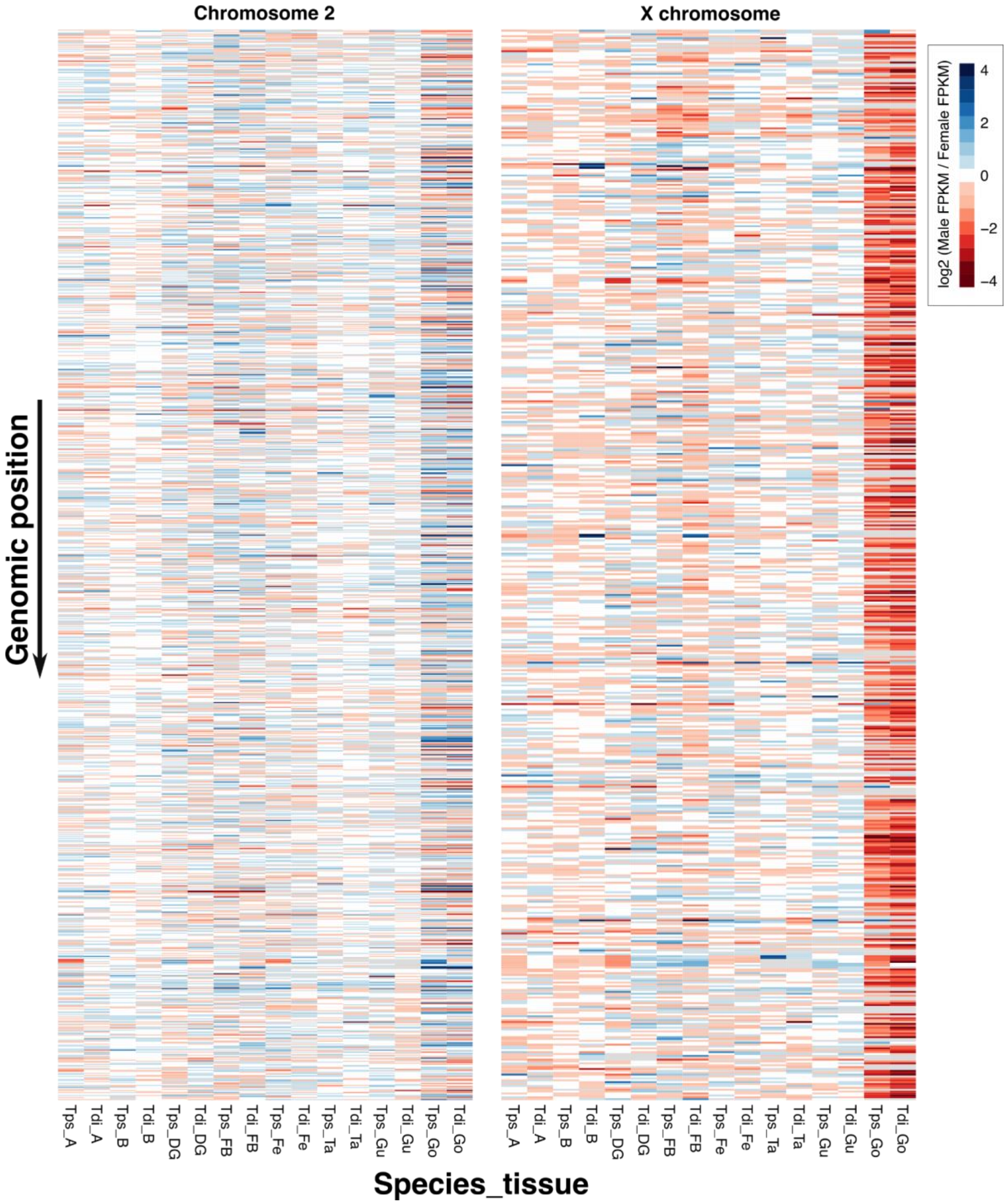


**Fig. S4 | Male to female expression ratios along the X and chromosome 3 for *T. poppense* (Tps) and *T. douglasi* (Tdi) in Antennae (A), Brain (B), Defence glands (DG), Fat body (FB), Femur (Fe), Gut (Gu), Tarsi (Ta), and Gonads (Go).**

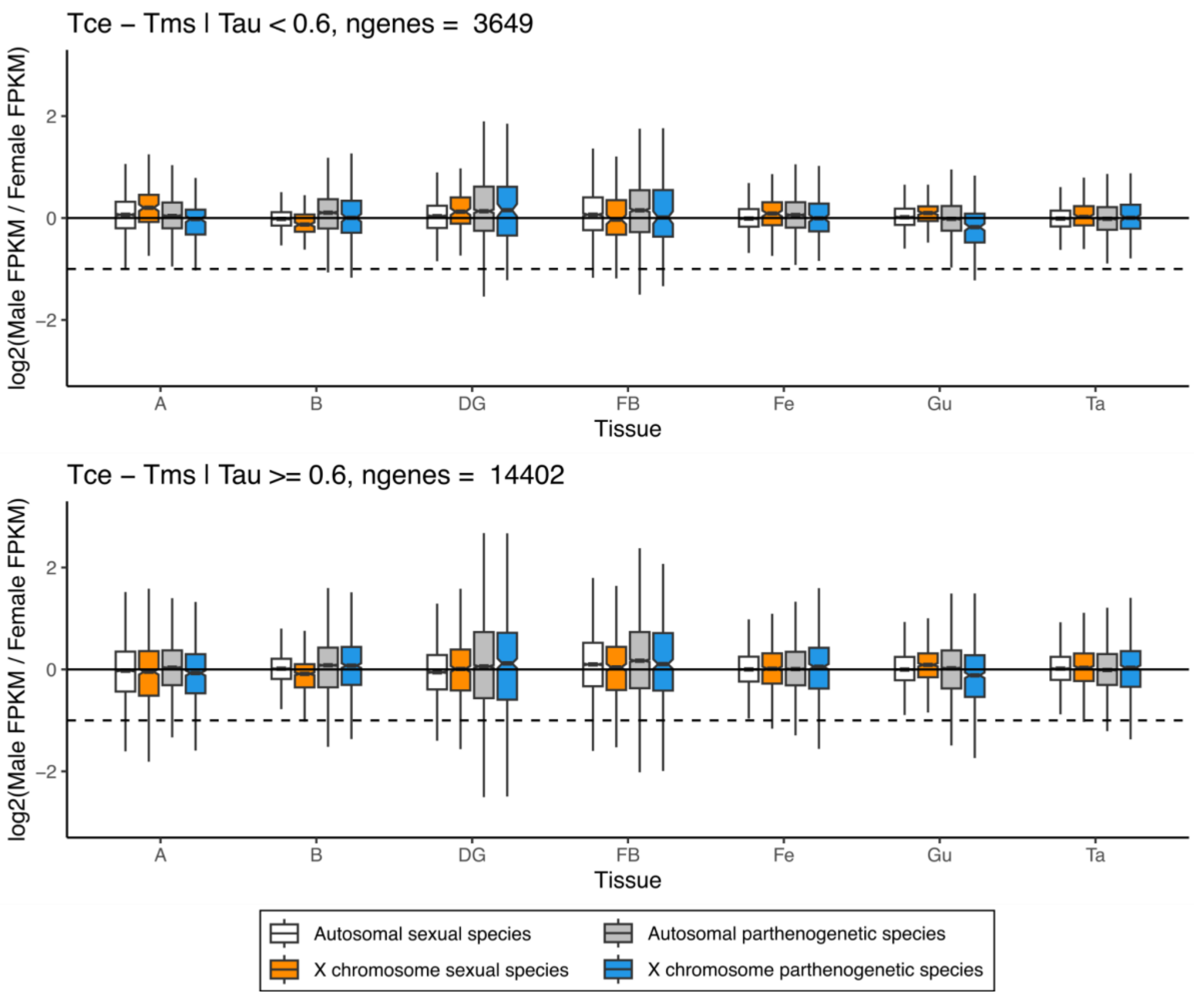


**Fig. S5 | Male to female expression ratios for the X and autosomes in non-reproductive tissues for genes with broad expression (Tau <0.6, top plot) and more tissue-specific expression (Tau >= 0.6, lower plot) in sexual species *T. cristinae* and parthenogenetic species *T. monikense*.** Dashed lines represent a two- fold reduction in expression in males (as expected if there was no dosage compensation).

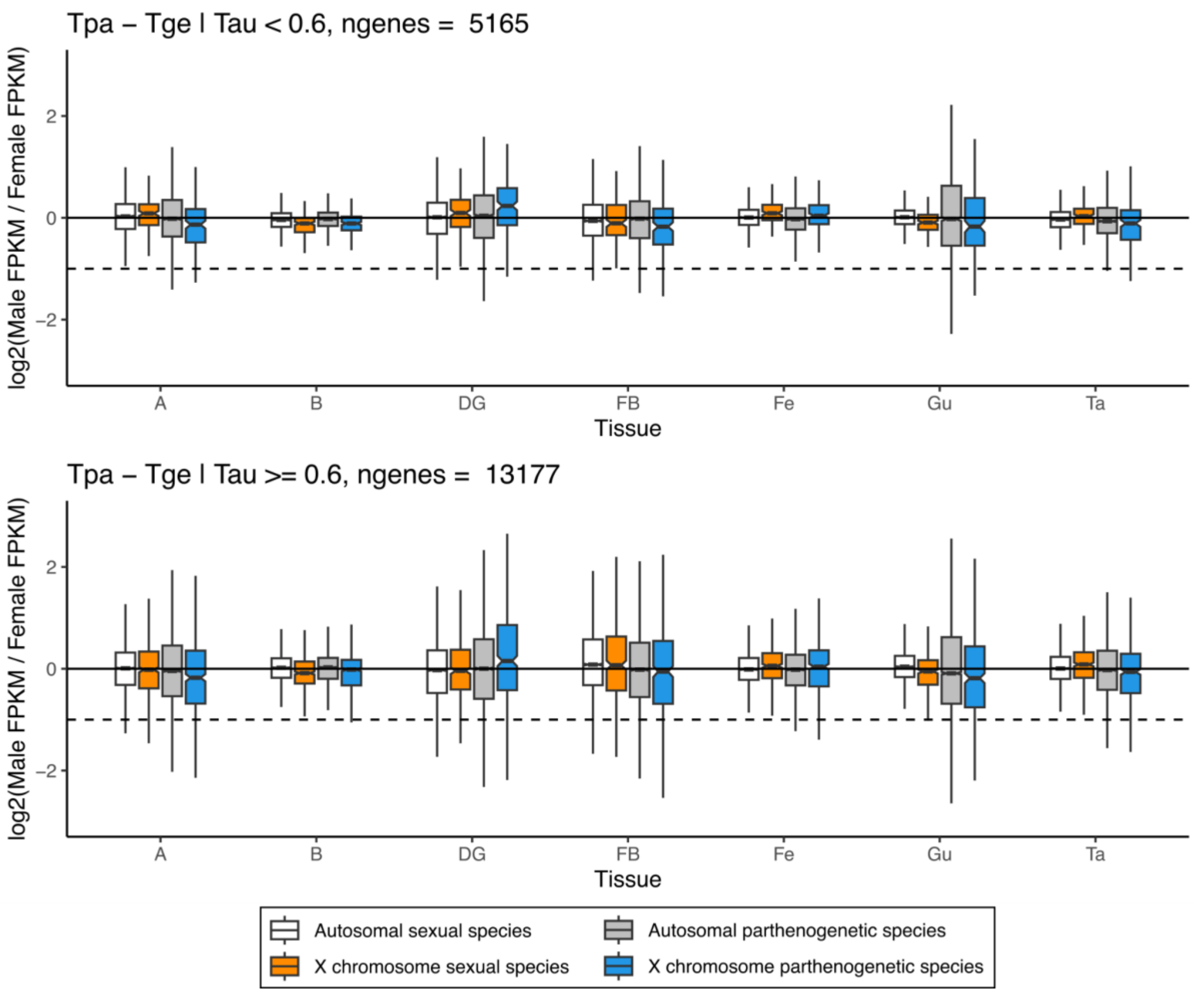


**Fig. S6 | Male to female expression ratios for the X and autosomes in non-reproductive tissues for genes with broad expression (Tau <0.6, top plot) and more tissue-specific expression (Tau >= 0.6, lower plot) in sexual species *T. podura* and parthenogenetic species *T. genevievae*.** Dashed lines represent a two- fold reduction in expression in males (as expected if there was no dosage compensation).

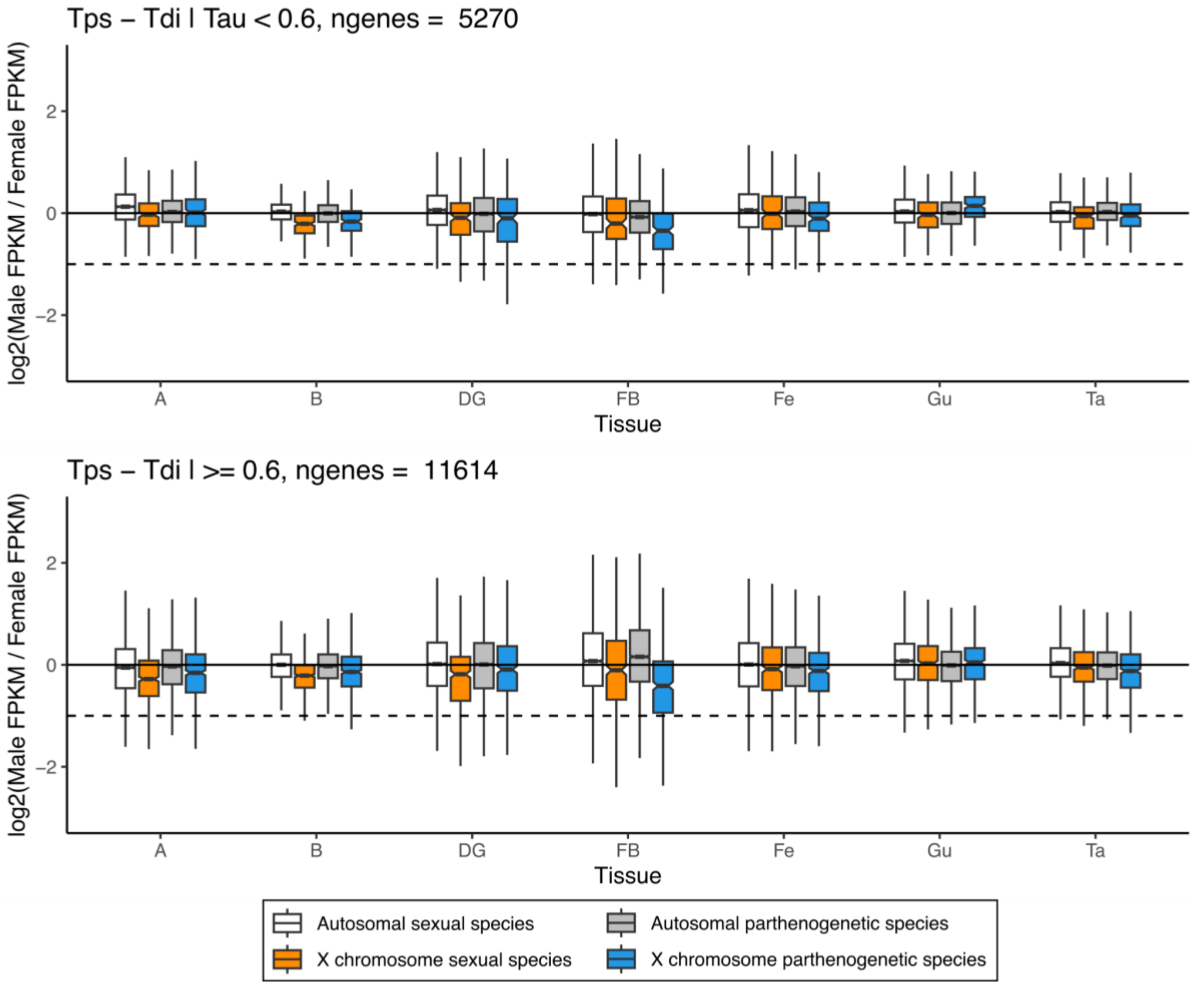


**Fig. S7 | Male to female expression ratios for the X and autosomes in non-reproductive tissues for genes with broad expression (Tau <0.6, top plot) and more tissue-specific expression (Tau >= 0.6, lower plot) in sexual species *T. poppense* and parthenogenetic species *T. douglasi*.** Dashed lines represent a two- fold reduction in expression in males (as expected if there was no dosage compensation).

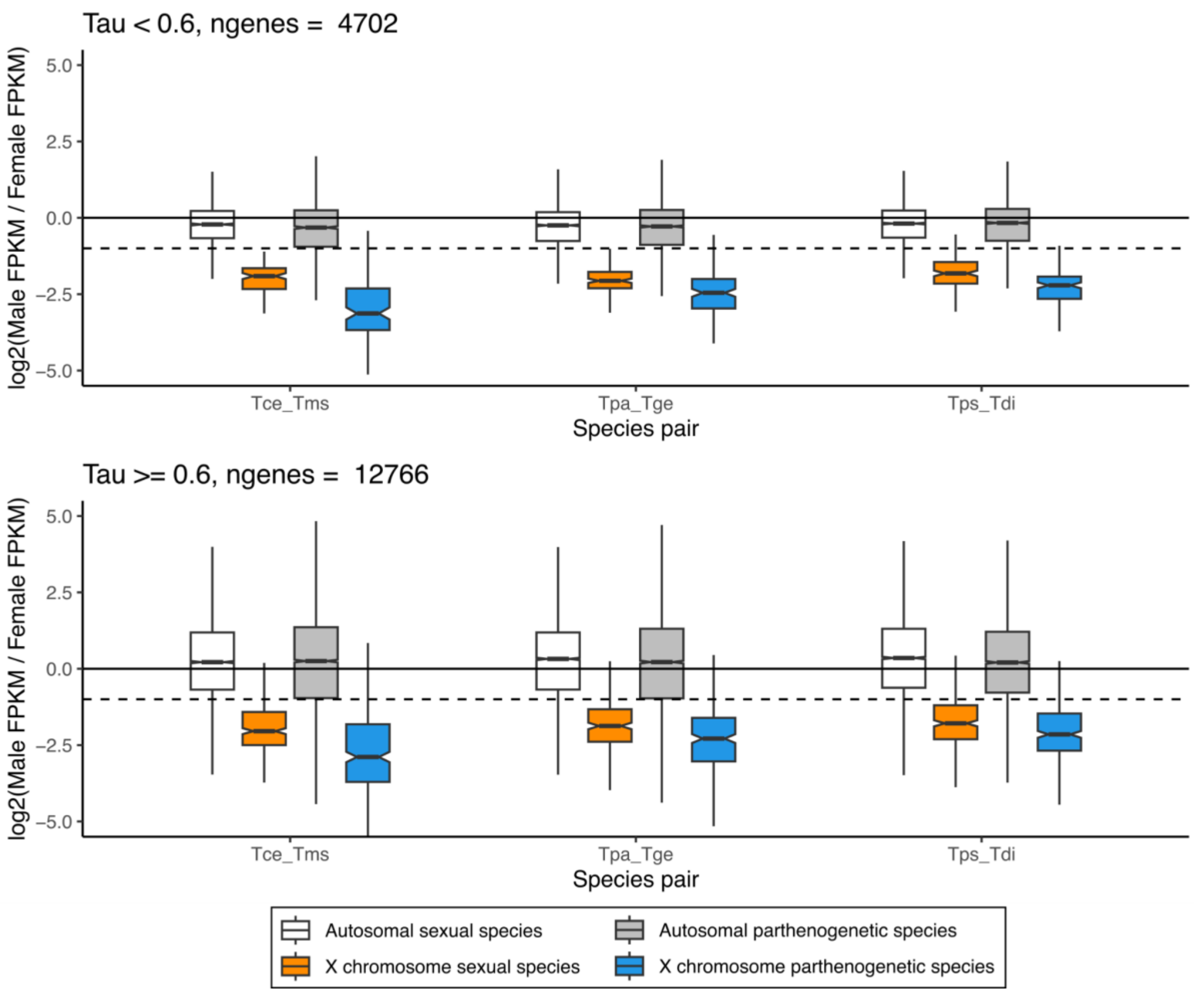


**Fig. S8 | Log$_2$ of male to female expression ratio for the X and autosomes in testes and female gonads for genes with broad expression (Tau <0.6, top plot) and more tissue-specific expression (Tau >= 0.6, lower plot). Species codes: Tce = *T. cristinae*, Tms = *T. monikense,* Tpa = *T. podura*, Tge = *T. genevievae*, Tps = *T. poppense*, Tdi = *T. douglasi.*** Dashed lines represent a two- fold reduction in expression in males (as expected if there was no dosage compensation).

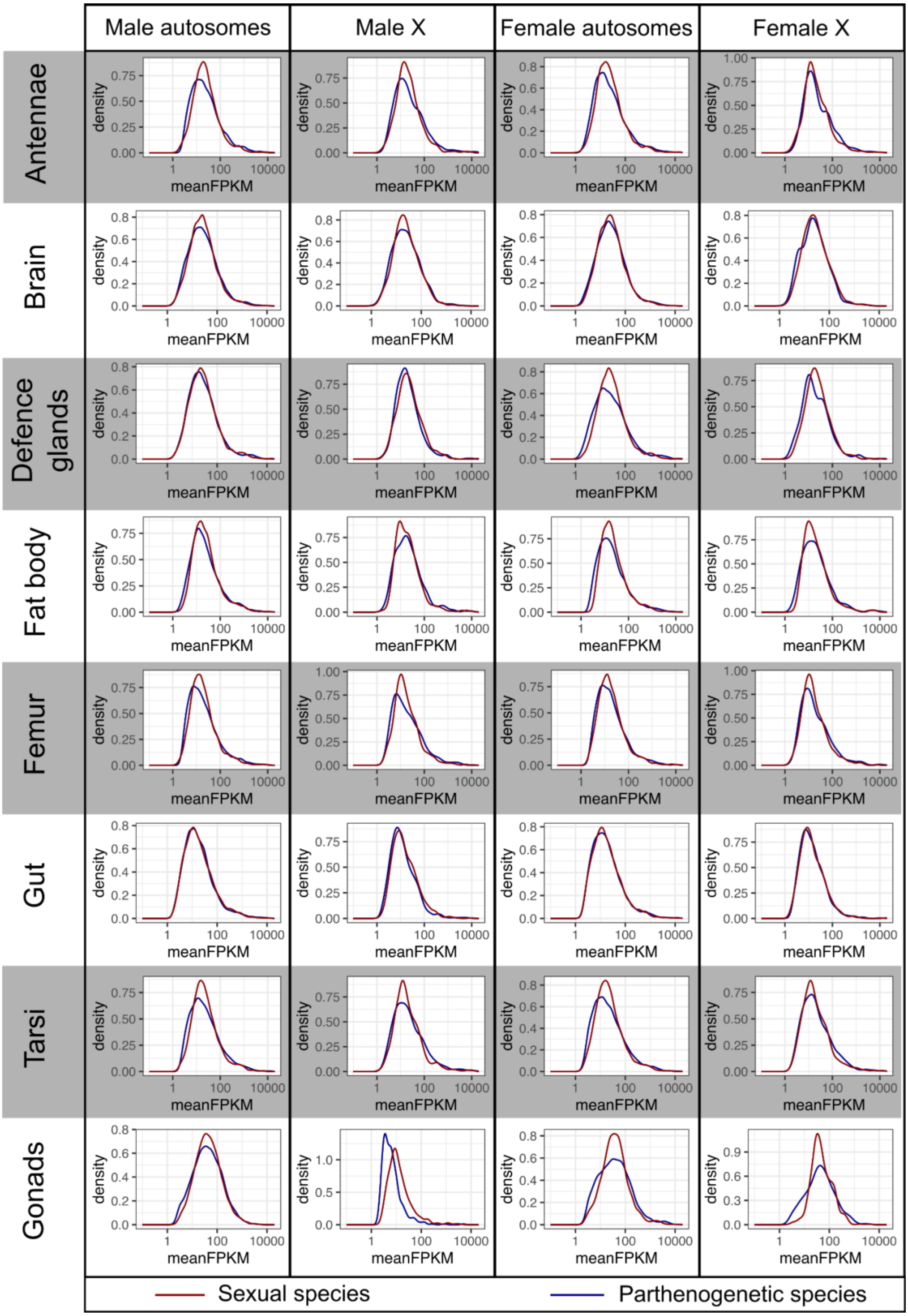


**Fig. S9 | Distribution of gene expression (FPKM) in sexual species *T. cristinae* and parthenogenetic species *T. monikense*.**

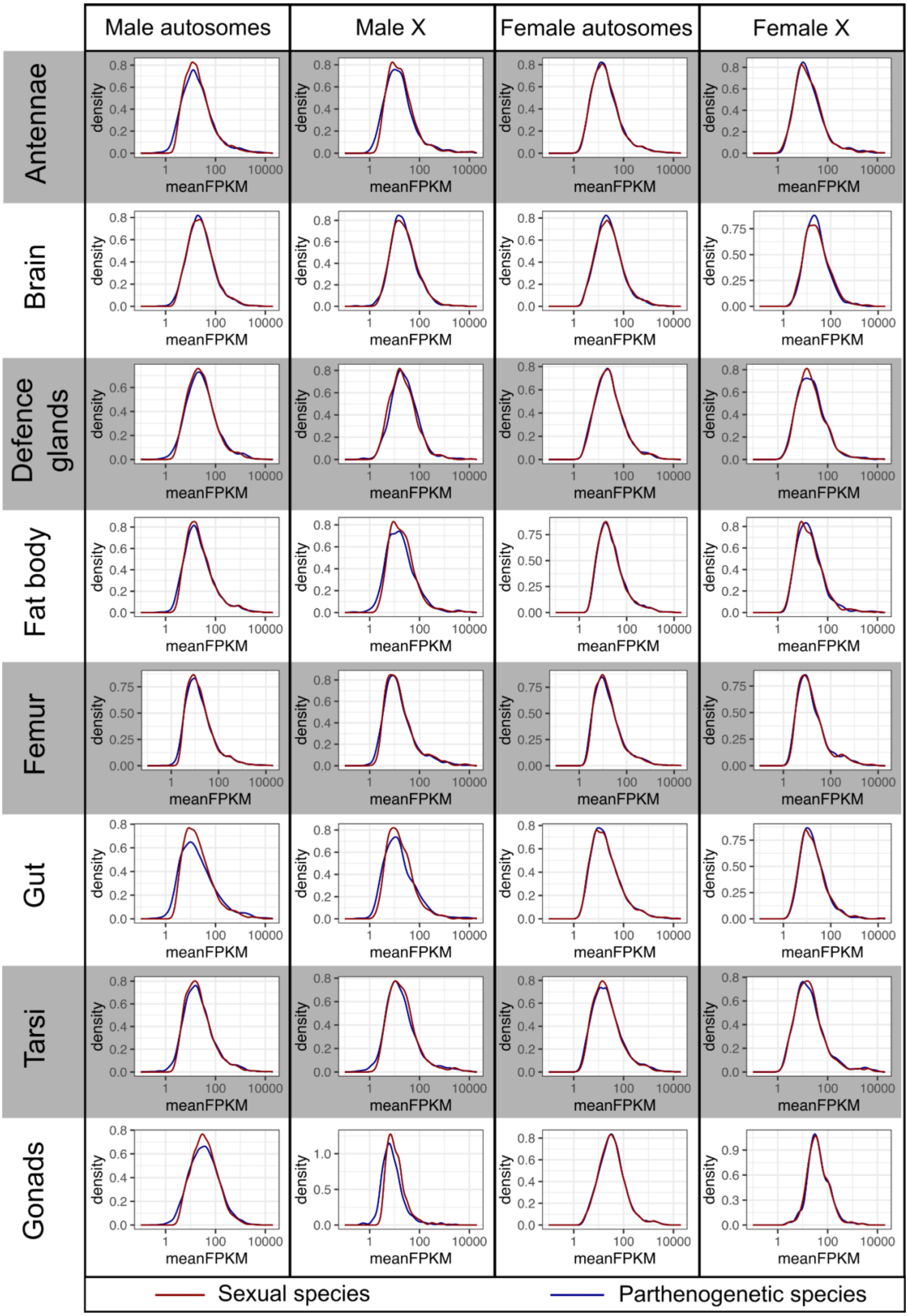


**Fig. S10 | Distribution of gene expression (FPKM) in sexual species *T. podura* and parthenogenetic species *T. genevievae*.**

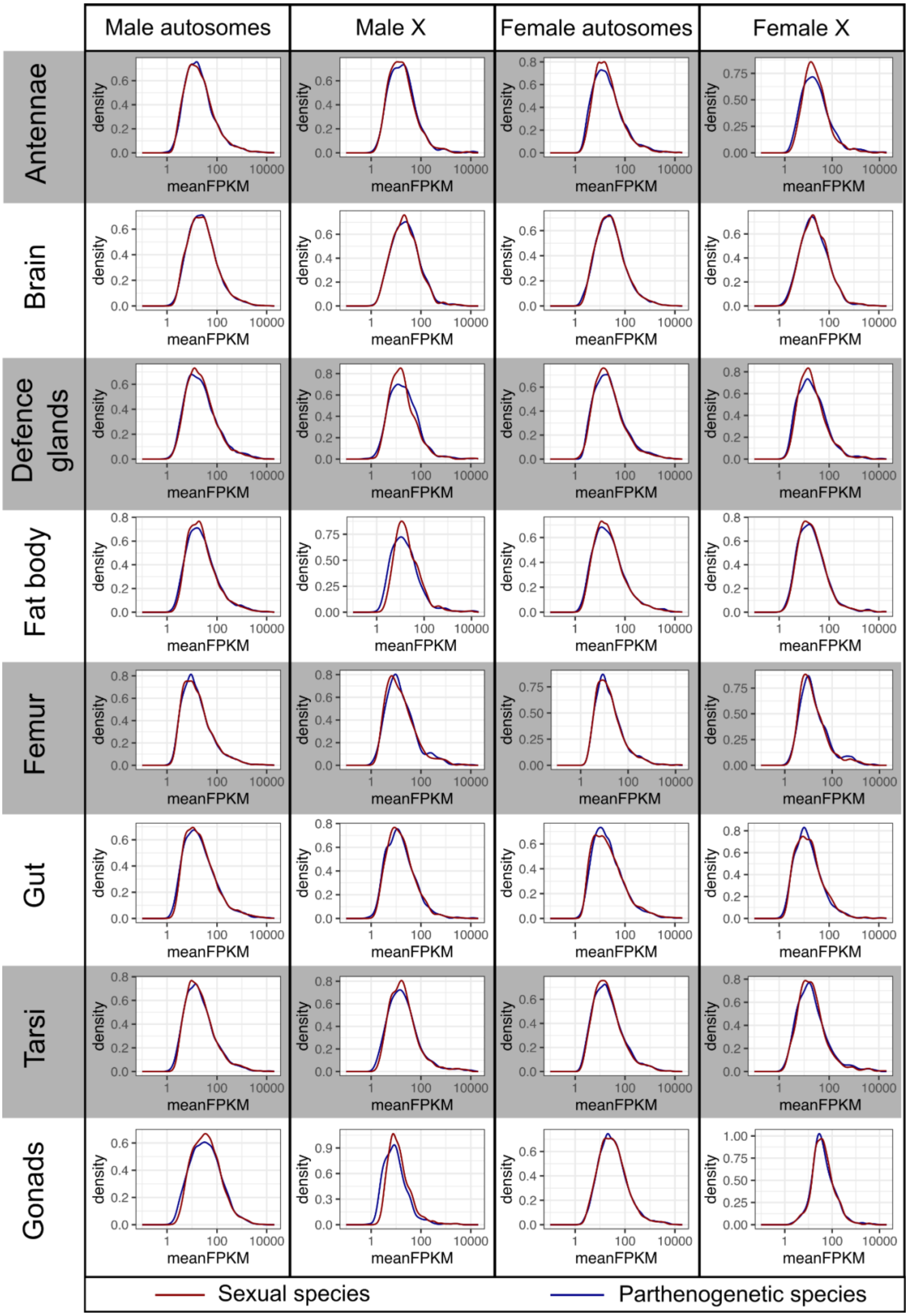


**Fig. S11 | Distribution of gene expression (FPKM) in sexual species *T. poppense* and parthenogenetic species *T. douglasi.***

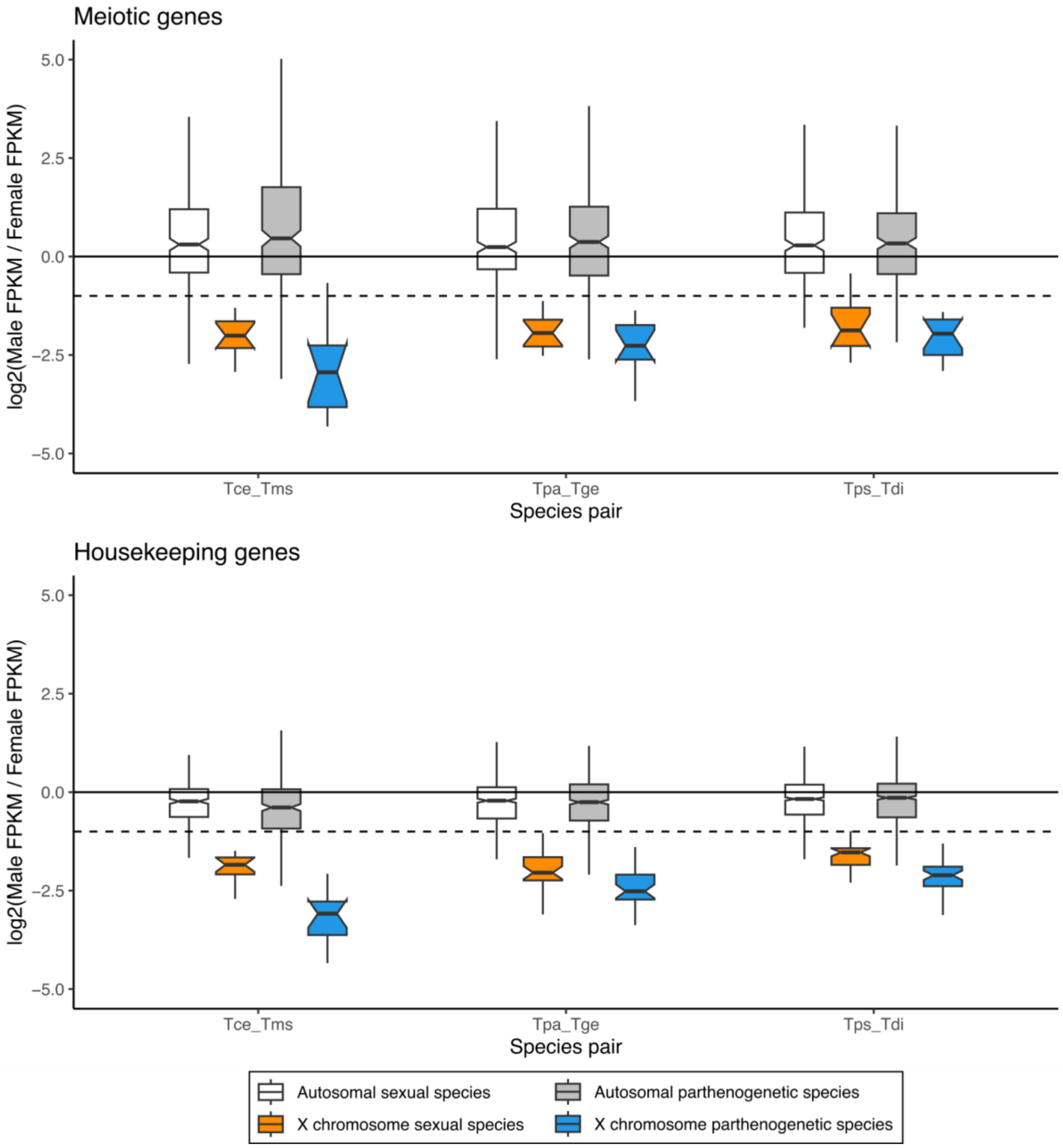


**Fig. S12 | $Log_2$ of male to female expression ratio for the X and autosomes in testes and female gonads for meiotic genes (genes annotated with the GO-term GO:0051321 (meiotic cell cycle) or one of its child terms, top plot) and housekeeping genes (genes showing consistent expression across the tissues (Tau <= 0.5), bottom plot).**

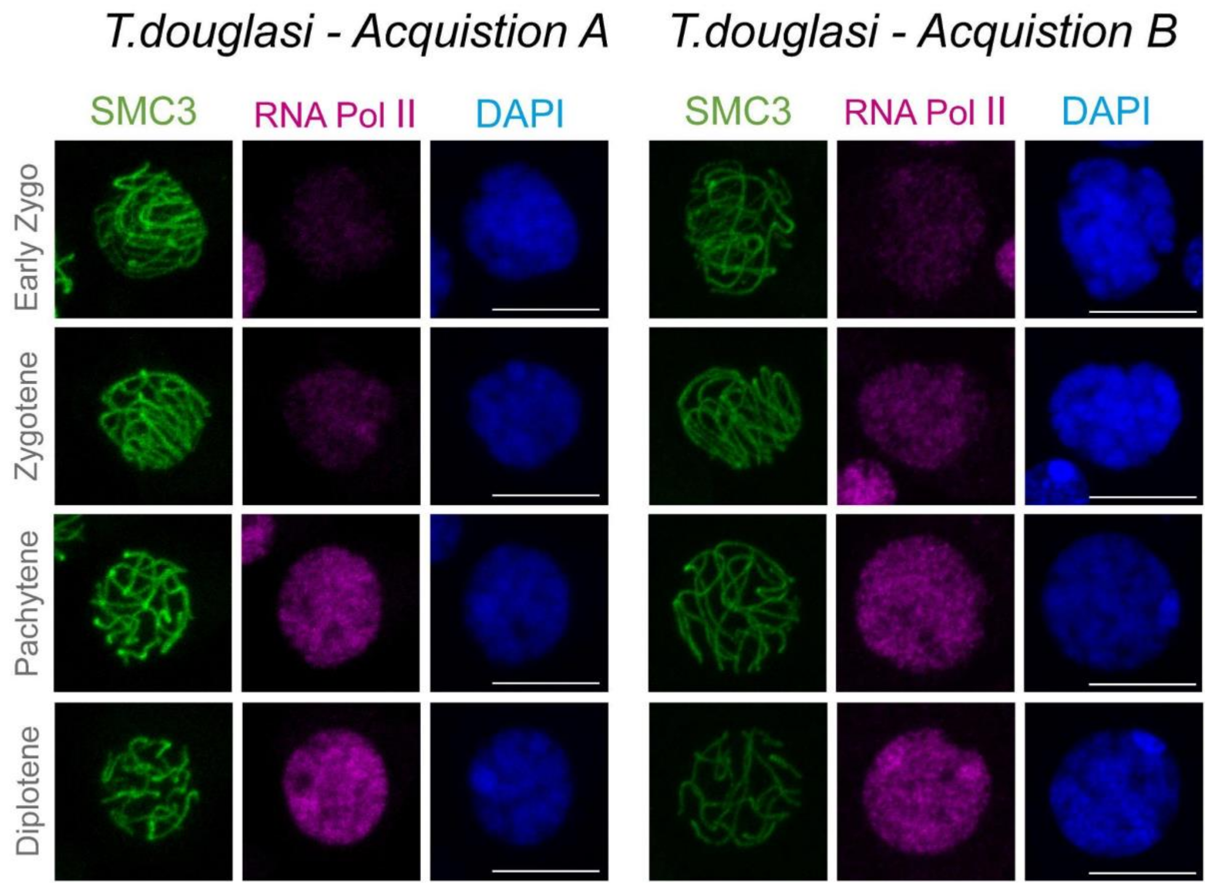


**Fig. S13 | Misregulation of autosomal gene expression in meiotic cells of parthenogenetic *T. douglasi* males.** RNApol-II distribution in *T. douglasi* meiocytes and spermatids. Projections of stack images acquired across *T. douglasi* squashed meiocytes , at the stages indicated, stained with DAPI (blue) and double immunolabeled for SMC3 (green) and RNApol-II (magenta). RNApol-II labelling is present already from early zygotene. Acquisitions from different individuals are shown to illustrate that the used immunolabeling allows for inference of qualitative (presence-absence) rather than quantitative (intensity of signals) differences (Fig. 4). Scale bar: 10 µm.

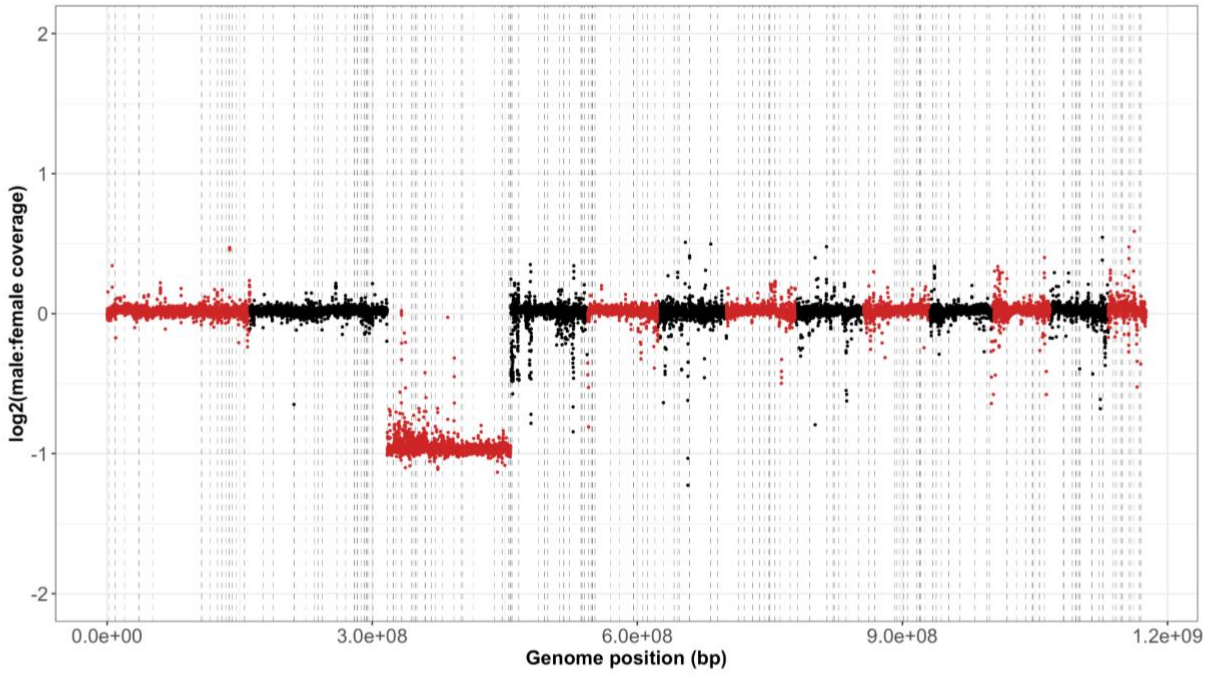


**Fig. S14 | Male to female coverage ratios for *T. cristinae.* Colours demark different linkage groups. Dashed lines represent breaks between scaffolds within a linkage group.**

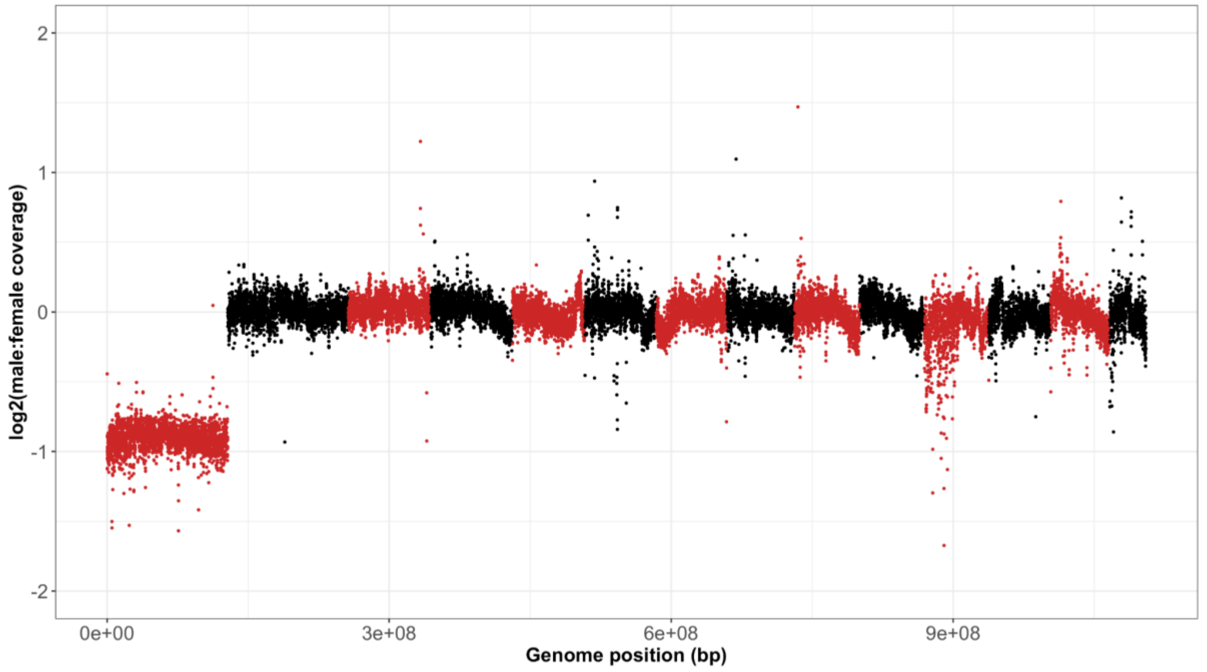


**Fig. S15 | Male to female coverage ratios for *T. podura.* Colours demark different linkage groups.**

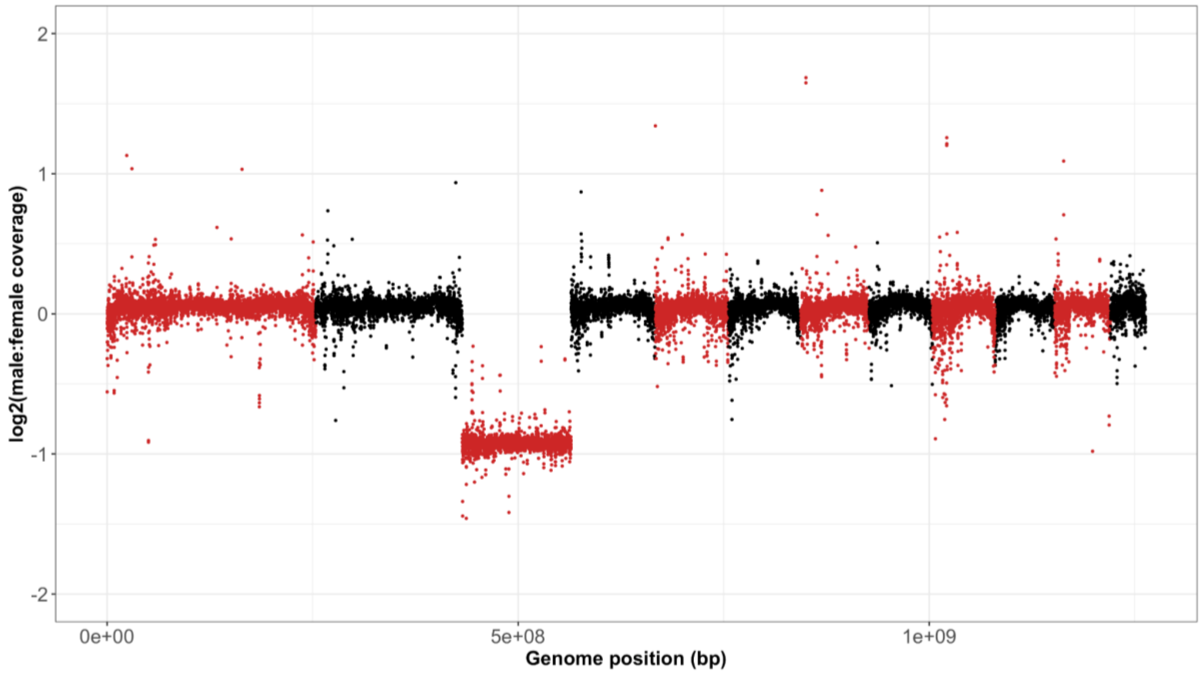


**Fig. S16 | Male to female coverage ratios for *T. poppense.* Colours demark different linkage groups.**

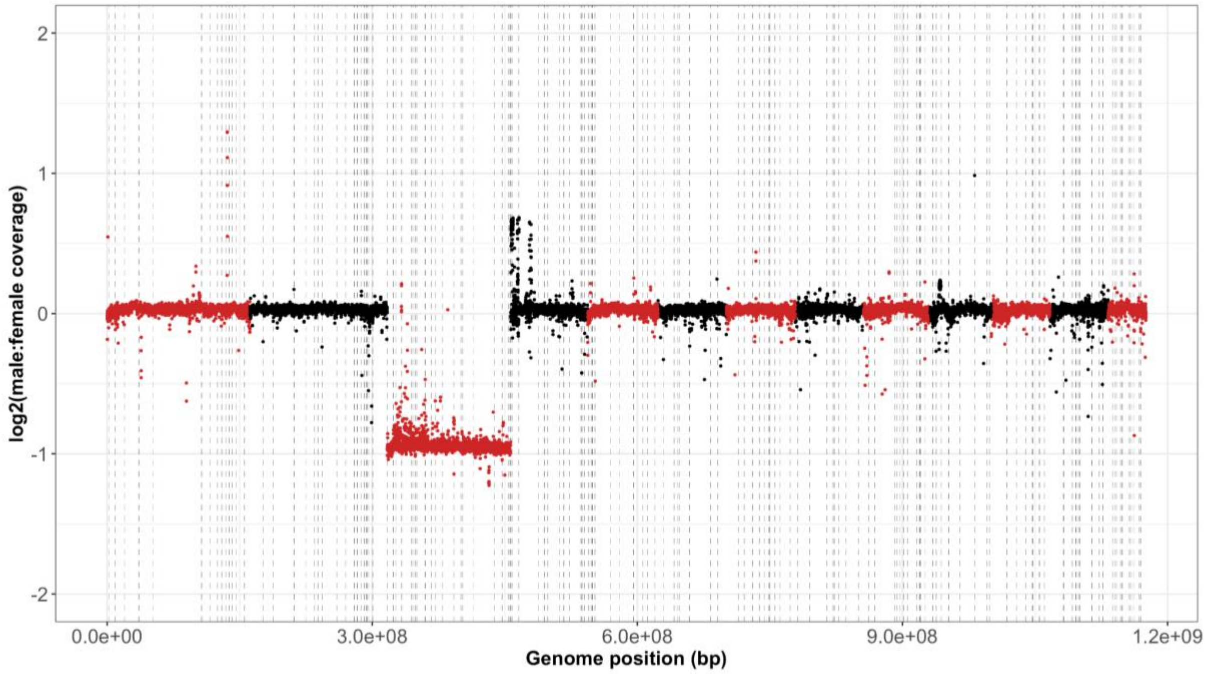


**Fig. S17 | Male to female coverage ratios for *T. monikense* mapped to the *T. cristinae* genome*.*** Colours demark different linkage groups. Dashed lines represent breaks between scaffolds within a linkage group.

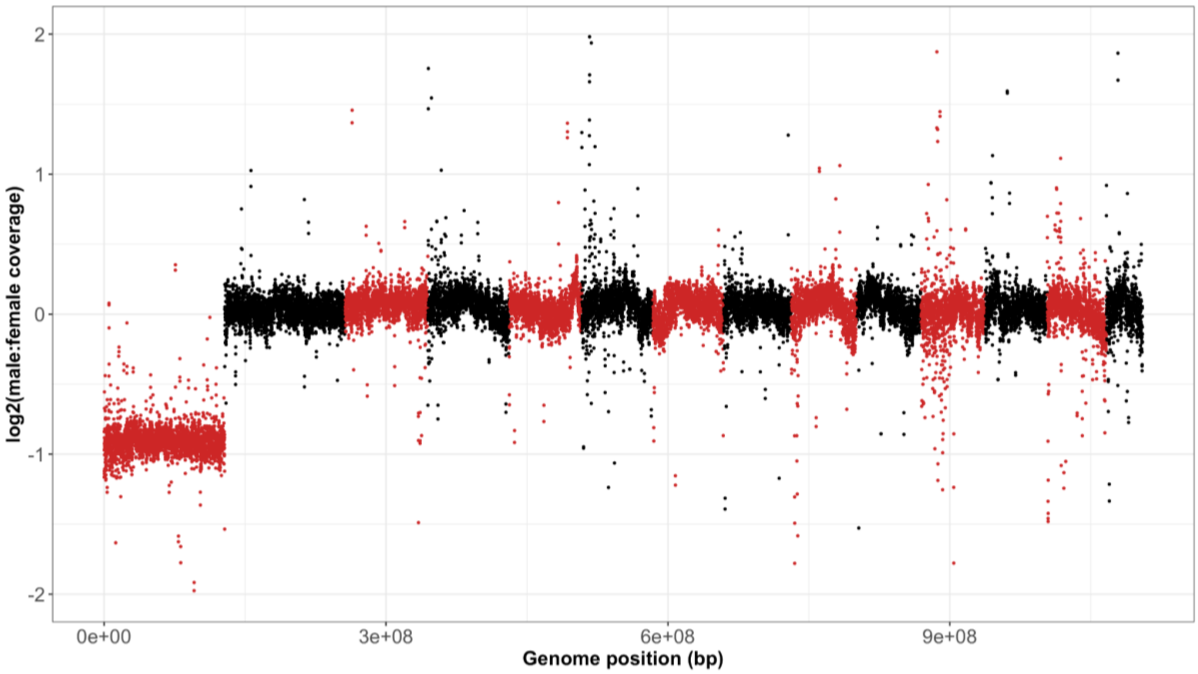


**Fig. S18 | Male to female coverage ratios for *T. genevievae* mapped to the *T. podura* genome*.*** Colours demark different linkage groups.

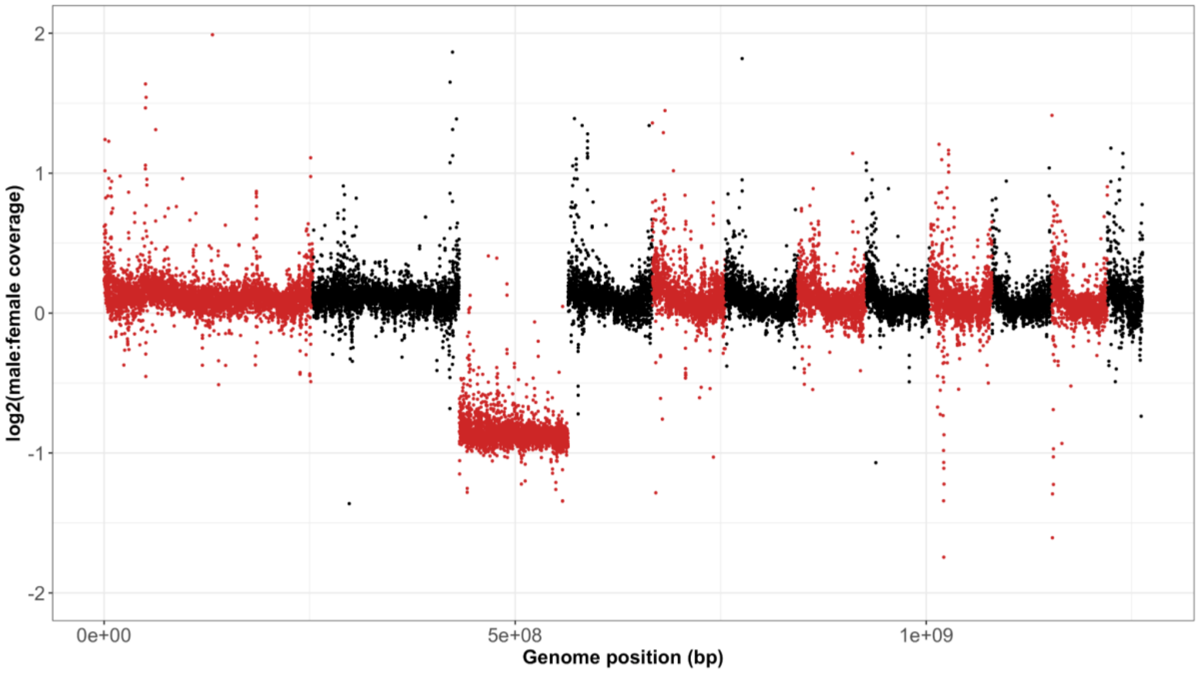


**Fig. S19 | Male to female coverage ratios for *T. douglasi* mapped to the *T. poppense* genome*.*** Colours demark different linkage groups.

### Supplementary tables

**Table S1 | Genome assembly statistics** Chromosome numbers indicate the haploid number in females inferred from karyotypes. Note that the data for *T. poppense* were generated in a previous study (17) and added here for comparability. *calculated on the un-scaffolded assembly. †calculated on the scaffolded assembly.

| | | | | | | Number of genes annotated | | |
|---|---|---|---|---|---|---|---|---|
| **Species** | **Size (Gb)** | **N50 (Mb)** | **n** | **% of assembly assigned to chromosomes** | **% BUSCO complete (fragmented)** | **Autosomes** | **X** | **Un-assigned** |
| *T. cristinae* | 1.21 | 10.1* | 13 | 95.1† | 97.3% (1.5) | 27759 | 2111 | 2002 (6.3%) |
| *T. podura* | 1.15 | 75.7 | 14 | 96.4 | 98.4% (0.7) | 29688 | 2262 | 1416 (4.2%) |
| *T. poppense* | 1.29 | 89.1 | 12 | 94.3 | 98.6% (1.0) | 34495 | 2148 | 3039 (8.1%) |

**Table S2 | Dosage compensation changes in non-reproductive tissues between sexual and parthenogenetic species.** delta MF (sexual) = Difference between $log_2$ of male to female expression ratio for the X and autosomes in sexual species. delta MF (parthenogenetic) = Difference between $log_2$ of male to female expression ratio for the X and autosomes in parthenogenetic species. The 'more unequal expression in' column indicates if the parthenogenetic or sexual species has more unequal expression between the X and autosomes for that tissue. Delta X - the difference in X expression between sexual and parthenogenetic species. FDR indicates the significance of Wilcoxon tests.

| Tissue | Species | delta MF (sexual) | FDR | delta MF (parth) | FDR | More unequal expression in | delta X | Higher X exp in | FDR |
|---|---|---|---|---|---|---|---|---|---|
| Antennae | Tce-Tms | -0.034 | 4.25E-01 | 0.093 | **4.86E-06** | Parth | 0.093 | Sexual | **3.86E-02** |
| Brain | Tce-Tms | 0.101 | **7.73E-20** | 0.028 | 9.68E-01 | Sexual | -0.153 | Parth | **2.68E-13** |
| Defence glands | Tce-Tms | -0.111 | **1.89E-03** | -0.051 | 9.99E-01 | Sexual | -0.056 | Parth | 1.74E-01 |
| Fat body | Tce-Tms | 0.070 | **4.75E-02** | 0.075 | 2.12E-01 | Parth | -0.070 | Parth | 1.05E-01 |
| Femur | Tce-Tms | -0.027 | 1.11E-01 | -0.017 | 9.99E-01 | Sexual | -0.014 | Parth | 7.19E-01 |
| Gut | Tce-Tms | -0.092 | **1.79E-07** | 0.159 | **1.37E-09** | Parth | 0.239 | Sexual | **5.50E-18** |
| Tarsi | Tce-Tms | -0.022 | 1.60E-01 | -0.036 | 2.97E-01 | Parth | 0.005 | Sexual | 7.13E-01 |
| Antennae | Tpa-Tge | -0.005 | 8.13E-01 | 0.123 | **4.70E-06** | Parth | 0.178 | Sexual | **1.12E-06** |
| Brain | Tpa-Tge | 0.096 | **1.24E-14** | 0.057 | **4.72E-07** | Sexual | -0.049 | Parth | 1.71E-01 |
| Defence glands | Tpa-Tge | -0.030 | 1.01E-01 | -0.170 | **5.71E-08** | Parth | -0.165 | Parth | **7.95E-06** |
| Fat body | Tpa-Tge | 0.011 | 4.12E-01 | 0.091 | **2.12E-02** | Parth | 0.124 | Sexual | **1.75E-03** |
| Femur | Tpa-Tge | -0.078 | **4.72E-07** | -0.062 | **8.05E-03** | Sexual | 0.029 | Sexual | 9.08E-02 |
| Gut | Tpa-Tge | 0.099 | **5.97E-14** | 0.106 | **7.16E-03** | Parth | 0.110 | Sexual | **1.86E-02** |
| Tarsi | Tpa-Tge | -0.081 | **3.90E-06** | 0.032 | **1.93E-02** | Sexual | 0.141 | Sexual | **1.88E-09** |
| Antennae | Tps-Tdi | 0.208 | **1.57E-19** | 0.101 | **4.17E-07** | Sexual | -0.083 | Parth | **6.21E-03** |
| Brain | Tps-Tdi | 0.220 | **1.73E-60** | 0.127 | **9.08E-16** | Sexual | -0.065 | Parth | **5.28E-05** |
| Defence glands | Tps-Tdi | 0.169 | **1.83E-14** | 0.096 | **2.70E-02** | Sexual | -0.036 | Parth | **1.40E-02** |
| Fat body | Tps-Tdi | 0.172 | **5.21E-07** | 0.441 | **3.76E-48** | Parth | 0.244 | Sexual | **3.56E-09** |
| Femur | Tps-Tdi | 0.092 | **1.45E-02** | 0.118 | **3.32E-05** | Parth | 0.052 | Sexual | 5.72E-02 |
| Gut | Tps-Tdi | 0.054 | **1.68E-02** | -0.089 | **4.63E-05** | Parth | -0.082 | Parth | **4.55E-02** |
| Tarsi | Tps-Tdi | 0.085 | **3.03E-07** | 0.099 | **1.26E-07** | Parth | 0.042 | Sexual | 7.82E-02 |

**Table S3 | Number of RNAseq samples used for annotation of reference genomes.** Species codes: Tce = *T. cristinae*, Tms = *T. monikense*, Tpa = *T. podura,* Tge = *T. genevievae.*

| Sex | Tissue | Stage | Tce | Tms | Tpa | Tge | Source |
|---|---|---|---|---|---|---|---|
| Female | Antennae | Adult | 4 | 0 | 0 | 0 | PRJNA504764 |
| Female | Antennae | Juvenile | 4 | 0 | 0 | 0 | PRJNA504764 |
| Female | Antennae | Adult | 4 | 8 | 4 | 4 | This study |
| Female | Brain | Adult | 4 | 5 | 5 | 4 | This study |
| Female | Defence glands | Adult | 5 | 4 | 4 | 4 | This study |
| Female | Fat body | Adult | 4 | 4 | 4 | 4 | This study |
| Female | Femur | Adult | 4 | 8 | 4 | 4 | This study |
| Female | Gonad | Adult | 4 | 5 | 4 | 4 | This study |
| Female | Gut | Adult | 4 | 4 | 4 | 4 | This study |
| Female | Tarsi | Adult | 4 | 8 | 4 | 4 | This study |
| Female | Whole-body | Hatching | 0 | 3 | 0 | 5 | PRJNA679785,PRJNA1295360 |
| Female | Whole-body | Adult | 6 | 3 | 6 | 3 | PRJNA380865 |
| Male | Accessory glands | Adult | 4 | 4 | 4 | 2 | This study |
| Male | Antennae | Adult | 4 | 0 | 0 | 0 | PRJNA504764 |
| Male | Antennae | Juvenile | 4 | 0 | 0 | 0 | PRJNA504764 |
| Male | Antennae | Adult | 4 | 7 | 4 | 2 | This study |
| Male | Brain | Adult | 4 | 4 | 4 | 2 | This study |
| Male | Defence glands | Adult | 4 | 4 | 4 | 2 | This study |
| Male | Fat body | Adult | 4 | 6 | 4 | 2 | This study |
| Male | Femur | Adult | 5 | 7 | 4 | 2 | This study |
| Male | Gonad | Adult | 4 | 7 | 4 | 2 | This study |
| Male | Gut | Adult | 4 | 4 | 4 | 2 | This study |
| Male | Tarsi | Adult | 4 | 4 | 4 | 2 | This study |
| Male | Whole-body | Adult | 3 | 0 | 3 | 0 | PRJNA380865 |
| Unknown | Whole-body | Hatching | 12 | 0 | 12 | 0 | PRJNA679785,PRJNA1295360 |

**Table S4 | Number of RNAseq samples used gene expression analyses.** Samples of *T. poppensis* and *T. douglasi* were obtained from our previous study (17). Species codes: Tce = *T. cristinae*, Tms = *T. monikense*, Tpa = *T. podura,* Tge = *T. genevievae,* Tps = *T. poppense* Tdi = *T. douglasi.*

| Sex | Tissue | Stage | Tce | Tms | Tpa | Tge | Tps | Tdi |
|---|---|---|---|---|---|---|---|---|
| Female | Antennae | Adult | 4 | 4 | 4 | 4 | 4 | 4 |
| Female | Brain | Adult | 4 | 4 | 4 | 4 | 4 | 4 |
| Female | Defence glands | Adult | 4 | 4 | 4 | 4 | 4 | 4 |
| Female | Fat body | Adult | 4 | 4 | 4 | 4 | 4 | 4 |
| Female | Femur | Adult | 4 | 4 | 4 | 4 | 4 | 4 |
| Female | Gonad | Adult | 4 | 4 | 4 | 4 | 4 | 4 |
| Female | Gut | Adult | 4 | 4 | 4 | 4 | 4 | 4 |
| Female | Tarsi | Adult | 4 | 4 | 4 | 4 | 4 | 4 |
| Male | Antennae | Adult | 4 | 3 | 4 | 2 | 4 | 3 |
| Male | Brain | Adult | 4 | 4 | 4 | 2 | 4 | 3 |
| Male | Defence glands | Adult | 4 | 4 | 4 | 2 | 4 | 3 |
| Male | Fat body | Adult | 4 | 3 | 4 | 2 | 4 | 3 |
| Male | Femur | Adult | 4 | 3 | 4 | 2 | 4 | 3 |
| Male | Gonad | Adult | 4 | 4 | 4 | 2 | 4 | 3 |
| Male | Gut | Adult | 4 | 4 | 4 | 2 | 4 | 3 |
| Male | Tarsi | Adult | 4 | 3 | 4 | 2 | 4 | 3 |

**Table S5 - Values of the autosomal and X male-to-female coverage ratio peaks for each species.** Species codes: Tce = *T. cristinae*, Tms = *T. monikense*, Tpa = *T. podura,* Tge = *T. genevievae,* Tps = *T. poppense* Tdi = *T. douglasi.*

| Species | Autosome peak | Sex_peak | Autosome len (bp) | X len (bp) | % X |
|---|---|---|---|---|---|
| Tce | 0.023 | -0.969 | 1035801681 | 139734914 | 11.9 |
| Tms | 0.035 | -0.958 | 1035801681 | 139734914 | 11.9 |
| Tpa | -0.010 | -0.915 | 976679347 | 128341569 | 11.6 |
| Tge | 0.050 | -0.885 | 976679347 | 128341569 | 11.6 |
| Tps | 0.070 | -0.923 | 1130673392 | 132376913 | 10.5 |
| Tdi | 0.090 | -0.885 | 1130673392 | 132376913 | 10.5 |

**Table S6 | Number of genes in each of the linkage groups in *T. cristinae.***

| Linkage group | N genes |
|---|---|
| 1 | 3803 |
| 2 | 3791 |
| 3 | 2111 |
| 4 | 2477 |
| 5 | 2384 |
| 6 | 1948 |
| 7 | 2194 |
| 8 | 2264 |
| 9 | 2179 |
| 10 | 2022 |
| 11 | 1786 |
| 12 | 1841 |
| 13 | 1070 |
| Not classified | 2002 |

**Table S7 | Number of genes in each of the chromosomes in *T. podura.***

| Chromosome | N genes |
|---|---|
| 1 | 2262 |
| 2 | 3268 |
| 3 | 2185 |
| 4 | 2638 |
| 5 | 2645 |
| 6 | 2371 |
| 7 | 2446 |
| 8 | 2117 |
| 9 | 2098 |
| 10 | 1958 |
| 11 | 2715 |
| 12 | 2094 |
| 13 | 2030 |
| 14 | 1123 |
| Not classified | 1416 |

**Table S8 | Number of genes in each of the linkage groups in *T. poppense.***

| Chromosome | N genes |
|---|---|
| 1 | 6314 |
| 2 | 4432 |
| 3 | 2148 |
| 4 | 3048 |
| 5 | 2730 |
| 6 | 2863 |
| 7 | 2716 |
| 8 | 2302 |
| 9 | 2186 |
| 10 | 2373 |
| 11 | 2193 |
| 12 | 1190 |
| Not classified | 3039 |
